# Mapping the Computational Similarity of Individual Neurons within Large-scale Ensemble Recordings using the SIMNETS analysis framework

**DOI:** 10.1101/463364

**Authors:** Jacqueline B. Hynes, David M. Brandman, Jonas B. Zimmermann, John P. Donoghue, Carlos E. Vargas-Irwin

## Abstract

The expansion of large-scale neural recording capabilities has provided new opportunities to examine multi-scale cortical network activity at single neuron resolution. At the same time, the growing scale and complexity of these datasets introduce new conceptual and technical challenges beyond what can be addressed using traditional analysis techniques. Here, we present SIMNETS, a mathematically rigorous and efficient unsupervised relational analysis framework designed to generate intuitive, low-dimensional neuron maps that support a multi-scale view of the computational similarity (CS) relations among individual neurons. The critical innovation is the use of a novel measure of computational similarity that is based on comparing the intrinsic structure of latent spaces representing the spiking output of individual neurons. We use three publicly available neural population test datasets from the visual, motor, and hippocampal CA1 brain regions to validate the SIMNETS framework and demonstrate how it can be used to identify putative subnetworks (i.e., clusters of neurons with similar computational properties). Our analysis pipeline includes a novel statistical test designed to evaluate the likelihood of detecting spurious neuron clusters to validate network structure results. The SIMNETS framework can facilitate linking computational geometry representations across scales, from single neurons to subnetworks, within large-scale neural recording data.

## 1.0 Introduction

The neural processes underpinning complex sensory, cognitive, and behavioral phenomena engage complex activation patterns across brain-wide networks (Fox et al., 2005; Greicius et al., 2003; Raichle et al., 2001; Sporns, 2013; Thomas Yeo et al., 2011). Within large-scale networks, smaller clusters of computationally interrelated neurons have been proposed to embody fundamental input-output processing modules responsible for perceptual integration, dexterous motor control, and memory storage or retrieval (Briggman et al., 2006; Harris, 2005; Truccolo et al., 2010; Yuste, 2015; Zagha et al., 2015). While recent technological advances have made it possible to record from ever larger neuronal populations at single neuron resolution (Buzsáki, 2004; Chung et al., 2022; Donoghue, 2002; Maynard et al., 1997; Paulk et al., 2022; Steinmetz et al., 2021, 2018), progress in understanding the computational organization of large networks has been hindered because of the lack of appropriate conceptual and technical frameworks that are mathematically principled, amenable to a range of statistical tools, flexible, scalable, and fast enough to work on large datasets (Brown et al., 2004; Gao and Ganguli, 2015; Paninski and Cunningham, 2018; Stevenson and Kording, 2011; Zador et al., 2022). Objectively identifying and characterizing the organizational features of subnetworks of neurons within and across brain areas would greatly simplify the process of tracking information flow within cortical circuits, modeling multi-scale neural dynamics, and understanding the general principles of neural information processing (Alivisatos et al., 2013; Kohn et al., 2020; Urai et al., 2022; Yuste, 2015).

How can computational subnetworks be identified? In cortical areas where the tuning properties of individual neurons are well-established, such as the primary visual (V1) and motor (M1) regions, the most straightforward approach to assessing computational inter-relationships would be to calculate tuning parameter similarities across neurons (Berman et al., 1987; Georgopoulos et al., 1982; Livingstone et al., 1996). However, simple parametric tuning models often fail to capture the temporal complexities of single neuron outputs and can perform poorly or even break down entirely under more ecologically or ethologically relevant experimental conditions (David et al., 2004; Hosman et al., 2021; Olshausen and Field, 2005; Pospisil and Bair, 2021). Moreover, classic single-neuron tuning function estimation techniques can miss or “average away” computationally-relevant features of individual spike trains (Churchland and Shenoy, 2007; Cunningham and Yu, 2014; Malik et al., 2011).

A variety of useful and important tools have been developed to study coordinated neuronal activity based on the underlying premise that similar spike patterns shared by a pair of neurons imply similar information processing properties (Abeles and Gat, 2001; Aertsen and Gerstein, 1985; Amarasingham et al., 2012; Gerstein et al., 1985; Gerstein and Michalski, 1981; Grün et al., 2002a; Kiani et al., 2015; Lopes-dos-Santos et al., 2013). These methods group neurons according to the similarity of spiking statistics (e.g., coincident spiking or correlated spike rate fluctuation). However, the mechanistic origin and computational significance of these correlations have proven more complex than initially theorized and are not fully understood (Brody, 1999; Cole et al., 2016; de la Rocha et al., 2007; Friston, 2011; Roudi et al., 2015; Smith and Kohn, 2008; Soudry et al., 2013). This has limited the utility and explanatory power of inter-neuronal correlations as a means for establishing the computational relatedness of neurons within a common feature space. These interpretational issues are further confounded by the significant statistical and computational challenges that come with implementing these methods at scale (Amarasingham et al., 2012; Brody, 1999; Cohen and Kohn, 2011; de la Rocha et al., 2007; Soudry et al., 2013).

Here, we present SIMNETS, an unsupervised relational analysis framework designed to generate low-dimensional neuron maps that quantify and support a multi-scale view of the computational similarity (CS) relations among individual neurons. For the purposes of our analysis, we define computation as the mapping of a set of inputs to a set of outputs according to a given set of rules (Agüera y Arcas et al., 2003; Kanerva, 2009). Under these conditions, the computational equivalence of any two neural systems should be sought not in the actual pattern of their spiking outputs *but in the relations of their outputs to one another within each system* (Kanerva, 2009; Shepard and Susan Chipman., 1970). Based on this principle, SIMNETS compares neurons based on the intrinsic relational structure of their firing patterns, represented by a spike train similarity (SSIM) matrix that captures the relative changes in activity across a set of predetermined time windows. In this way, each SSIM matrix can be considered to represent the “output space” of a neuron, which serves as a computational “fingerprint” within the context of a given dataset (Penido et al., n.d.; Vargas-Irwin et al., 2015a). Using a SSIM matrix representation of single neuron output spaces make it possible to quantify the computational similarity relationships among a population of simultaneously recorded single units in a computationally efficient way using relatively simple metrics (e.g., matrix correlation or regression). As we will demonstrate, this 2nd-order spike train comparison makes it possible to identify neurons’ with computationally similar output spaces even when they use different encoding schemes: for example, if one neuron responds to a particular set of (unknown) conditions with an increased firing rate, and another responds to the same set of conditions with a stereotyped, precisely timed pattern of spikes, their respective SSIM matrices will display high similarity values in the same entries (even though spike trains each neuron emits are quite different). The Computational Similarity (CS) of the neurons is the single neuron SSIM matrices are represented in a pairwise Computational Similarity (CS) matrix, which can then be projected into a low dimensional map such that each point represents a neuron and the distance between them corresponds to their degree of computational similarity. The geometry of this neuron embedding can facilitate the identification of discrete clusters of neurons with distinct functions, as well as more complex shapes and gradients. This approach makes it possible to efficiently scale to large numbers of trials and neurons, making the method particularly suited for naturalistic settings with large numbers of possibly encoded features to be assessed.

## 2.0 Methods

The flow chart presented in Figure 1 provides a general overview of the main steps used to analyze neural ensemble data using the SIMNETS framework:

1. *Selecting spike trains.* Time series data representing the activity of N simultaneously recorded neurons (e.g., N spike times sequences) are split into S equal duration time segments that correspond to experimentally relevant events of interest.
2. *Generating a Spike train Similarity (SSIM) matrix for each neuron.* Use an appropriate metric (e.g., VP edit-distance)(Victor and Purpura, 1996) to describe the intrinsic pairwise similarities among each neuron’s set of S spike trains.
3. *Calculating the NxN Computational Similarity (CS) matrix.* Calculate the pairwise similarities (e.g., Pearson’s *r*) among all pairs of single neuron SSIM matrices. The resulting set of CS scores is represented as a population CS matrix, where each entry depicts the similarities among the output space of a neuron pair and depicts the overall pattern of relationships among the neurons for the experiment.
4. Visualizing results using dimensionality reduction (DR). An appropriate DR method (e.g., PCA-initialized t-SNE) can be applied to the SSIM matrices generated in step 2 to produce SSIM maps where each point represents an individual spike train, and the distance between them represents their relative similarity (Hinton and van der Maaten, 2008). This step makes it easier to interpret the neuron’s relationship to task variables (e.g., stimulus inputs, movement types, outlier trials) and ultimately their computational role within the context of higher-level network structures. Applying DR to the CS matrix generated in step 3 will produce a low-dimensional CS map, with each point corresponding to an individual neuron, and the distance between them corresponds to their relative computational similarity (i.e., the similarity between their respective SSIM matrices).

**Fig. 1.**
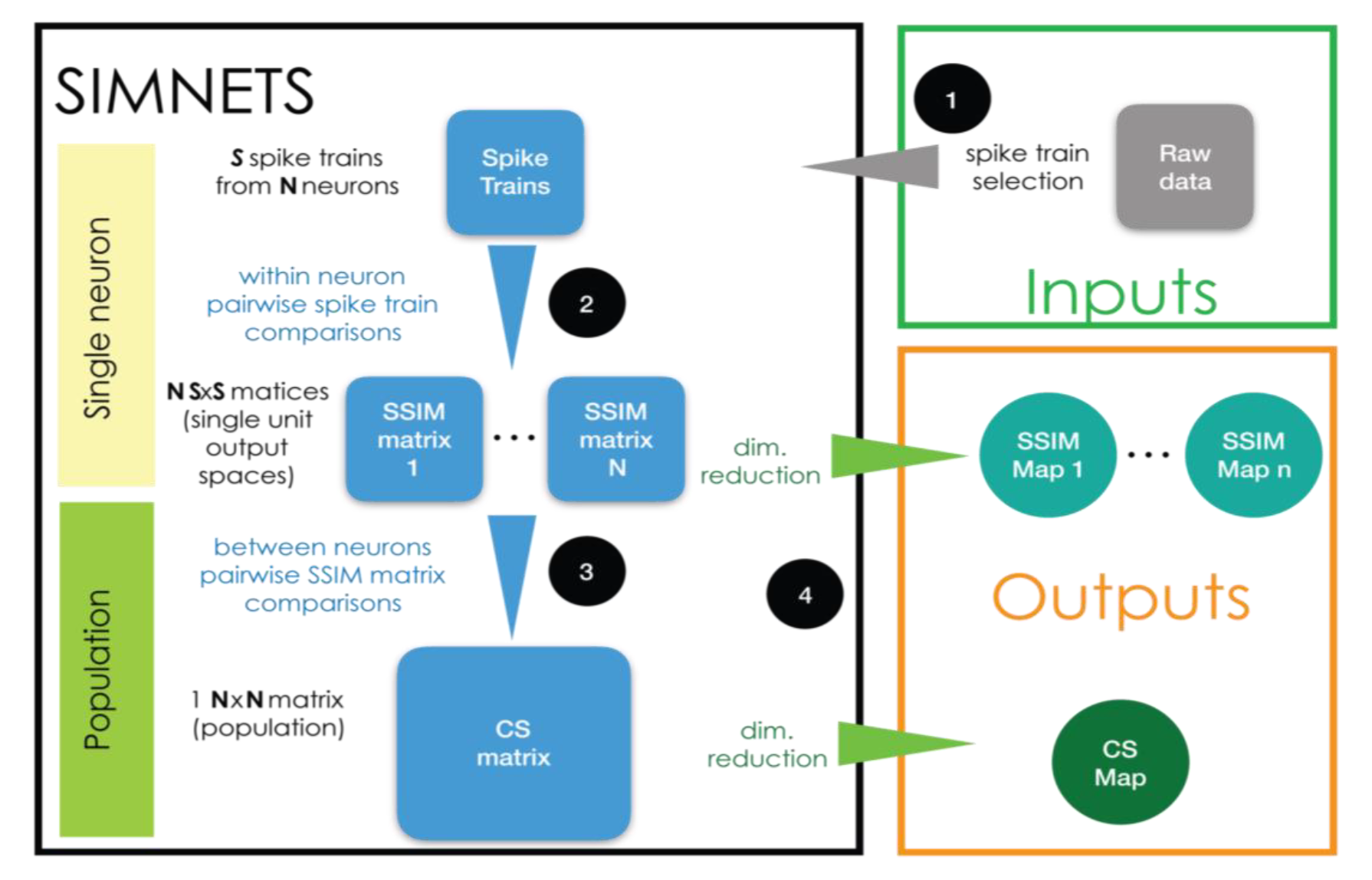
Flow diagram of the SIMNETS analysis framework.

The CS map is the main output of the algorithm, capturing the computational organization of the ensemble and serving as a starting point for further analysis aimed at identifying groups of neurons with similar computational properties. The following sections and figures describe specific steps (Fig. 2 – 3) and subsequent analysis (Fig. 4) in more detail.

**Fig. 2.**
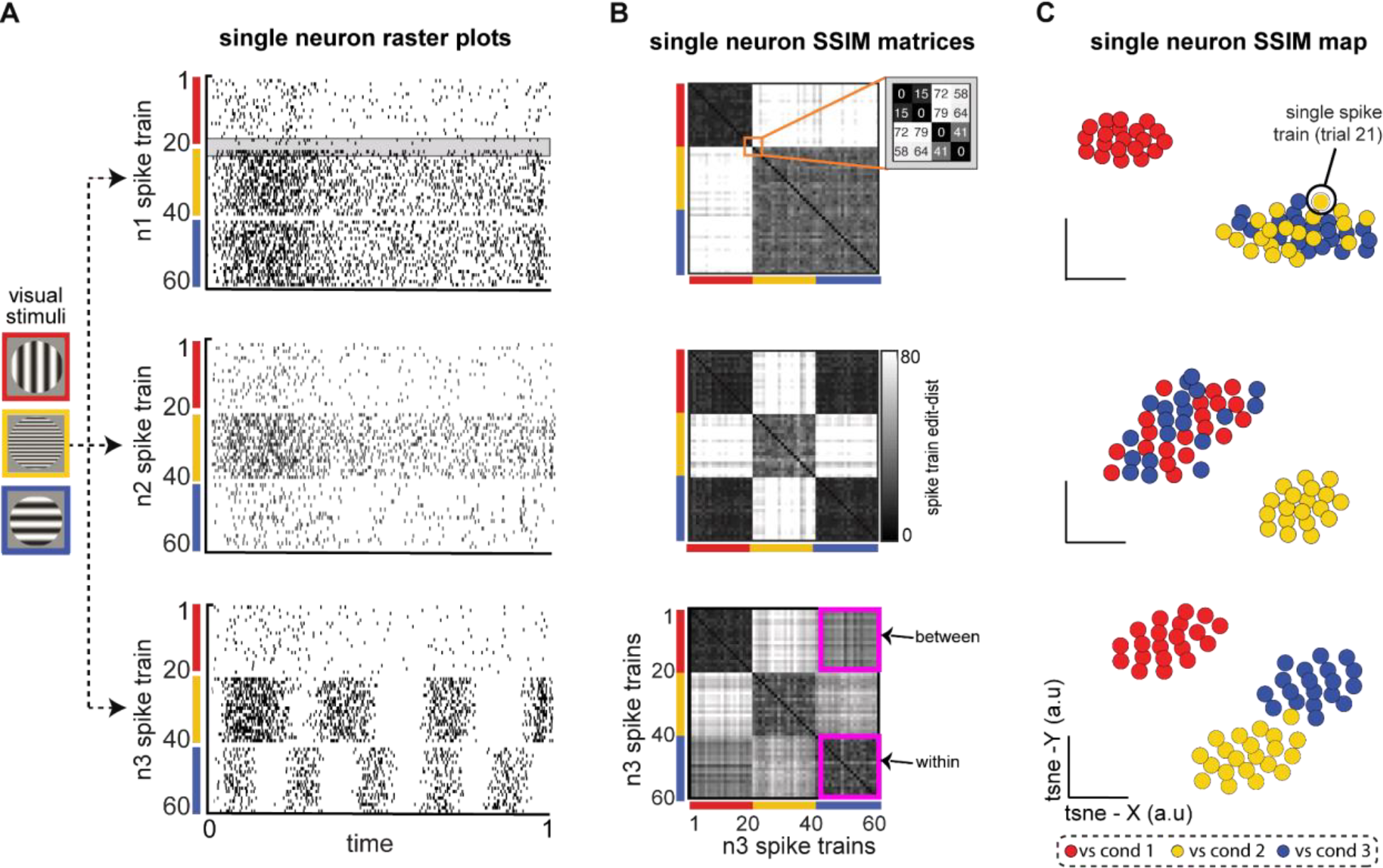
Single neuron Spike train Similarity (SSIM) matrices depict the computational geometry of each neuron’s spike train output space (OS) without the need for extrinsic labels or tuning models. **A)** Raster plots for three computationally distinct LNP model V1 neurons (n1, n2, n3) during simulated receptive field stimulation (Meyer et al., 2016). Visual stimulus (VS) conditions included vertically oriented grating of low spatial frequency (SF) (red, cond 1); horizontally oriented and high SF grating (yellow, cond 2); and horizontally orientated and low SF grating (blue, cond 3). Gray shaded region (n1 raster plot) highlights a similar pair of spike trains from VS condition 1 (S19 – S20) and a similar pair from VS condition 2 (S21 – S22). **B)** Neuron n1 (top), n2 (middle), n3 (bottom) single neuron SSIM matrices depict the SxS pairwise similarities among a neuron’s spike trains (a within-neuron comparison). Similarities are described in terms of “cost-based” edit distance (grayscale color bar), where smaller distance values correspond to similar spike trains (black, lowest edit-cost), and increasingly large distance values are increasingly dissimilar spike train outputs (white, larger edit-cost). The orange square in the top row (n1 SSIM matrix) highlights the set of distance values for the example spike trains S19 – S22 (highlighted in **(A)**). The magenta squares in the (bottom row; n3 single neuron SSIM matrix) highlight the matrix region that contains containing “within-condition” edit-distances (n3 cond-1, n3 cond-1) and “between-condition” distances (n3 cond-1, n3 cond-3). **C)** Three single neuron SSIM maps, the low-dimensional projection of each neuron’s single neuron SSIM matrix (from **(B)**). The intrinsic geometry of each neuron’s spike train output space provides a richer representation of single-neuron functional responses than standard parametric or statistical descriptions of single neuron responses (e.g., trial-averaged tuning curves).

**Fig. 3.**
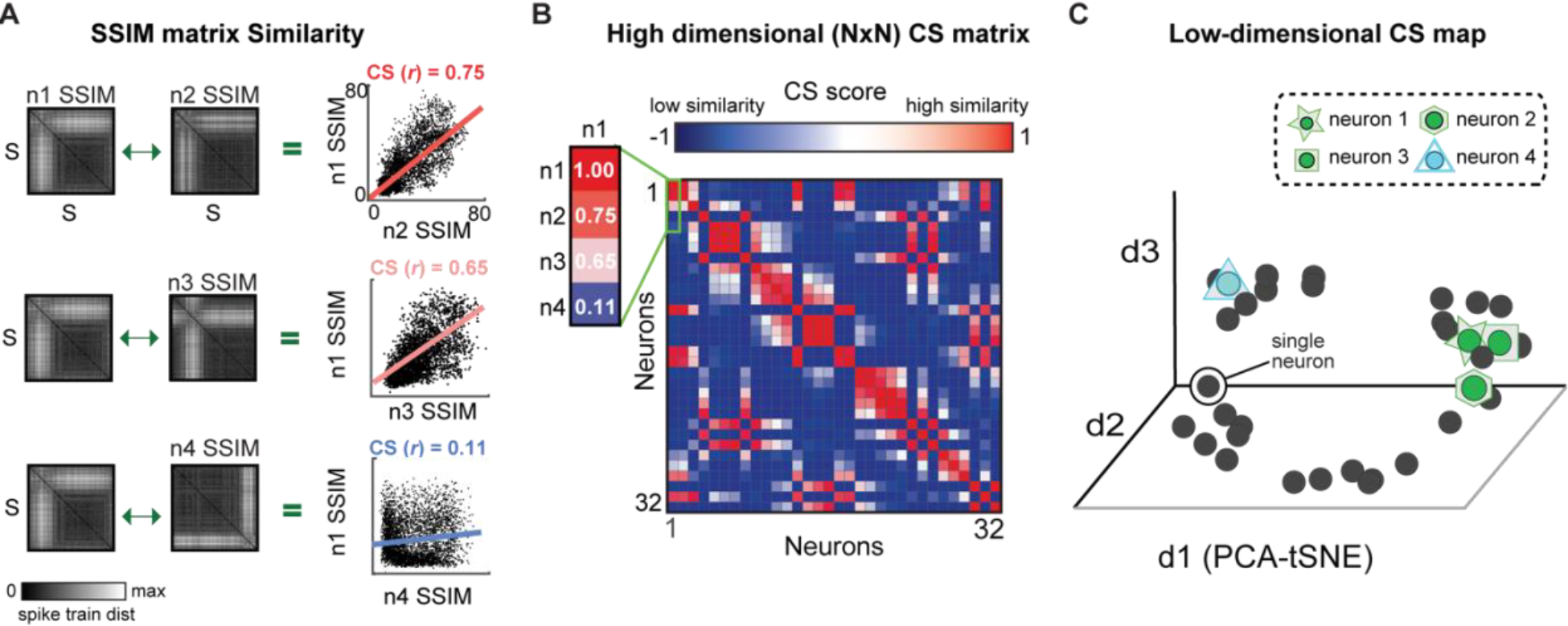
The similarity between the output spaces of all individual neurons is represented as a computational similarity (CS) matrix. **A)** Computational similarity is calculated by correlating each pair of SSIM matrices, illustrated for three neuron pairs. The regression line depicts the relationship among the VP distance values of a neuron pair. The visual similarity of the top two matrices (n1 v n2) and the differences in the bottom two (n1 v n4) is captured in their respective correlation values. **B)** Population CS matrix, highlighting the correlation values shown in A. **C)** CS Map of neuron similarity across the population obtained by applying DR to the CS matrix. Each cluster of neurons reflects a group neuron with similar computational properties.

**Fig. 4.**
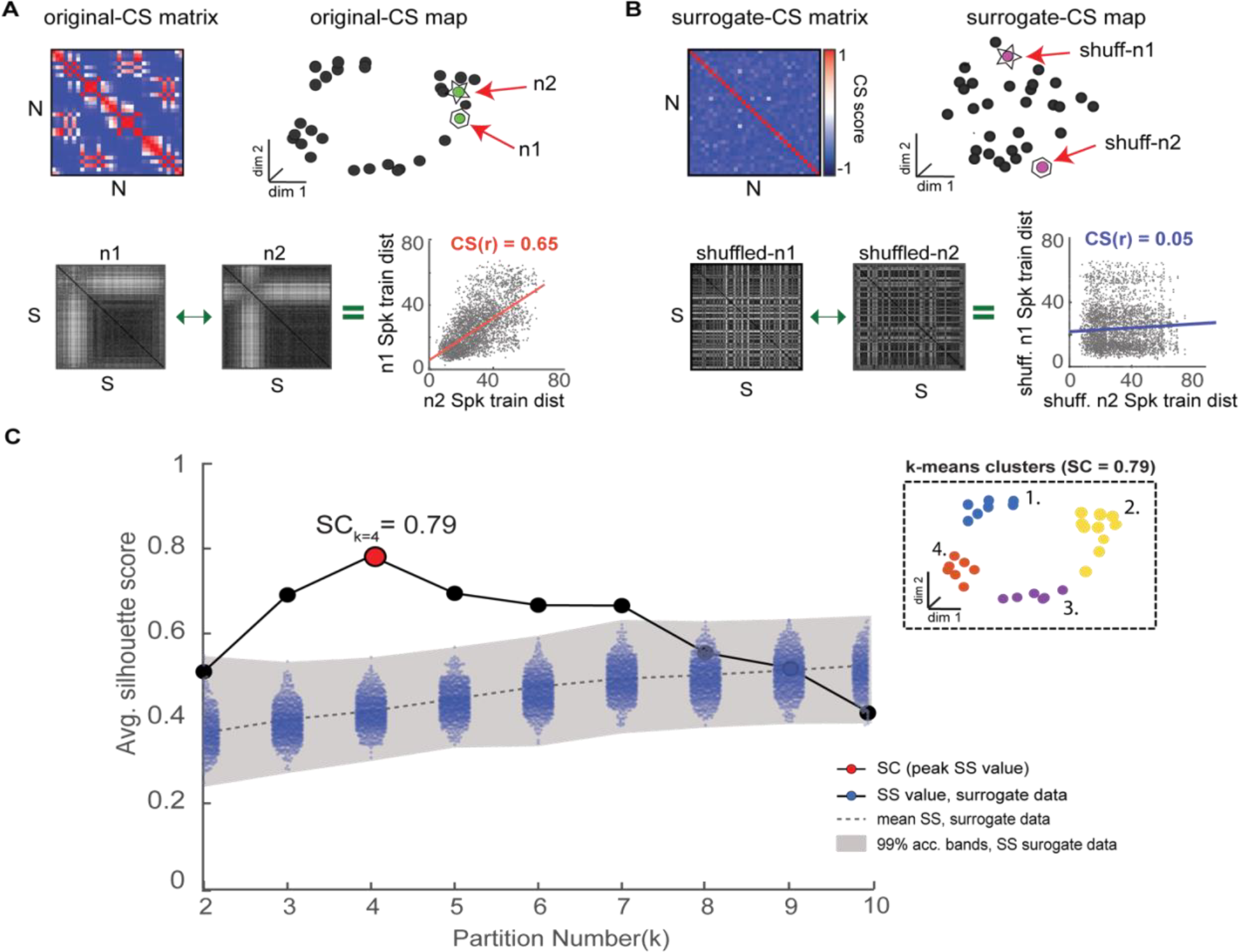
Novel shuffle-based silhouette analysis is used to detect and validate CS neuron clusters within an observed CS neuron map. The goal is to determine if an observed CS neuron map contains statistically meaningful neuron clusters. **A)** Top: population CS matrix and map for an example simulated neuron population (same as Fig. 3). Bottom: Two example single neuron SSIM matrices (n1 and n2) for a neuron pair with a high CS score (CS(r) = 0.65). **B)** Top: Surrogate CS matrix and map generated using CS neuron scores obtained after independently randomizing spike train order in all single neuron SSIM matrices. Bottom: SSIM matrices for the neurons shown in A after shuffling display reduced CS scores. **C)** The shuffling procedure is repeated many times (e.g.,10000) to generate an empirical chance distribution of silhouette values and obtain acceptance bands. Shading denotes the 99% confidence interval of surrogate data, which serves as an acceptance band for the null hypothesis (see (Amarasingham et al., 2012; Schemper, 1984) for further discussion). The silhouette values of the original data (without shuffling) are overlaid in black. The highest silhouette value (highlighted in red. is used to select the optimal number of clusters. Clusters corresponding to the highest (Ksc = 4) are shown in the inset.

### Representing single neuron output spaces using SSIM matrices

One of the main innovations of the SIMNETS framework consists of using single neuron SSIM matrices as a “computational fingerprint” to represent the intrinsic structure of the output space of individual neurons. This is accomplished by performing within-neuron measures of pairwise spike train similarity across all spike trains for each neuron, resulting in an SxS single neuron SSIM matrix for each of the N neurons (Fig. 1 step 2, Fig. 2B). Multiple approaches to estimate the similarity between pairs of spike trains have been proposed (Paiva et al., 2010) and could be used for this step. Here, we use the spike train metric proposed by Victor and Purpura (Houghton and Victor, 2010; Victor and Purpura, 1998, 1996)(See Box 1 for description).

### Box 1: Victor and Purpura Edit-based Spike Train Metric

This VP metric is a type of edit-distance for quantifying differences between pairs of spike trains and computed as the total “cost” of transforming one spike train into another through a series of elementary operations (Victor, 2005; Victor and Purpura, 1996).

These elementary operations include (1) inserting a spike, (2) deleting a spike, and (3) shifting a spike in time. Inserting or deleting a spike has a cost of c = 1 and shifting a single spike in time has a cost proportional to the amount of time that it is moved (c = qΔ t). The set of edits-steps associated with the minimum total edit-cost defines the shortest path between two points (spike trains) in the neuron’s spike train metric-space. The q parameter influences the relative importance of spike count and spike time differences when assessing spike train similarities. When q = 0, the cost of shifting a spike to the desired location will always be cheaper than deleting and re-inserting a spike in a spike train. Thus, for D(q=0), the minimum cost is simply the difference in the number of spikes between the spike trains. As the q value is increased beyond zero, spike timing begins to impact the cost of matching the spike trains. In this way, q controls the temporal resolution of the spike train comparison. In the context of the SIMNETS algorithm, a high q parameter will bias the algorithm toward grouping neurons based on information encoded over fine timescales, whereas a low q parameter will bias the algorithm toward grouping neurons based on the information encoded over coarse timescales; the temporal accuracy of the algorithm can be characterized as 1/q. The VP method has the advantage of operating in a point process framework, allowing for comparisons between relatively long spike trains (on the order of seconds) while preserving details of millisecond scale spike timing, by changing the cost assigned to shifting spikes in time (q parameter in the VP algorithm, see supplemental methods for details). Additionally, a point-process metric space can capture non-linearities in the neuron’s output space (Aronov and Victor, 2004; Fernandez and Farell, 2009).

In the current work, each single neuron SSIM matrix represents the relationship (i.e., similarity) between all spike trains for a single neuron. Fig. 2. presents three examples of SSIM matrices derived from the spike trains of simulated neuron responses (Linear Non-Linear Poisson functional model) (Meyer et al., 2016). Entries equal to zero (black) in the matrices represent identical spike trains (zero VP cost), while higher values represent increasing differences between the spike trains. For ease of viewing, the rows/columns of the matrices have been sorted according to simulated experimental conditions. Note that a specific ordering of the spike trains is not required for the subsequent comparisons (and is only included for illustrative purposes). It can be useful to graphically illustrate the relationship between individual spike trains across the experiment captured by the single neuron SSIM matrices using dimensionality reduction (Fig. 2B). Here, we use PCA-initialized tSNE (Hinton and van der Maaten, 2008): a combined unsupervised dimensionality reduction technique that aims to preserve the local neighborhood structure, via t-SNE, as well as information related to the global shape, via PCA-initialization (Kobak and Linderman, 2021; Lee et al., 2015). The non-random PCA-initialization also ensures reproducibility across iterations (Hinton and van der Maaten, 2008). This mapping step is not necessary for the following steps of the SIMNETS analysis pipeline, but these maps can be useful to visualize and compare the features coded in the population. For example, in Fig. 2B, the single neuron SSIM maps illustrate the separation of orientation selectivity in simulated neuron n1, the spatial frequency selectivity in n2, and the combined spatial and orientation selectivity of n3 (*See* Fig. S1 and “Supplementary Information” for additional discussion and examples). Changes in the shape of the SSIM output space map may also indicate the influence of other variables which are not explicitly manipulated.

## Computational Similarity Map: Comparing Single Neuron Output Spaces

Each SSIM matrix characterizes the intrinsic geometry of the output space for a specific neuron. The next step of the algorithm compares these “computational fingerprints” across the full set of recorded neurons. This second-order comparison is made by correlating each pair of SSIM matrices (Fig. 3; Fig. 1, step 3). Here we use Pearson’s correlation (Fig. 3A). Each correlation value is entered into a symmetrical (N x N) matrix (Fig. 3B) representing the relationships across neurons, which we term the Computational Similarity (CS) matrix. Applying dimensionality reduction (PCA-initialized t-SNE, (Hinton and van der Maaten, 2008)) to the CS matrix creates a CS map (Fig. 3C; Fig. 1, step 4). Note that, while each colored point in a single neuron SSIM map represents an individual spike train (Fig. 2C), each point in the CS map represents a neuron (Fig. 3C). In the population CS maps, the distance between neurons is a relative measure of the similarity of computational fingerprints to other neurons in the population. Therefore, neurons performing similar computations will tend to aggregate in clusters within the population CS map of neurons.

## Identifying potential subnetworks: unsupervised cluster detection & validation procedure

After neurons are embedded in the CS map, the task of finding groups of neurons representing potential neuronal subnetworks can be addressed using a wide variety of clustering algorithms. Here, we use the k-means clustering algorithm, an efficient centroid-based clustering method, however, other density-based, distribution-based, or hierarchical-clustering methods would also be appropriate. The k-means algorithm works by iteratively assigning neurons into the pre-specified number k clusters until the optimal solution is reached. However, the challenge is to ensure that the clustering structures identified by the k-means algorithm reflect genuine clusters within the CS neuron map and not false discoveries. We combined the k-means algorithm with two different cluster validation techniques, a silhouette graphical analysis and our novel shuffle-based statistical test, which together help avoid false cluster discovery (Fig. 4).

Silhouette analysis is a validation technique used to determine the most likely number of clusters in a dataset (Fig. 4) (Rousseeuw, 1987). A silhouette value represents the ratio of the *between* to *within-cluster* distances: a point within an ideal cluster will be close to members of the same cluster and far from points assigned to different clusters, resulting in a high silhouette value. Choosing the cluster number with the highest average silhouette value, the Silhouette Coefficient (SC) score, maximizes the separation between potential clusters (Fig. 4C, red point). We use a novel shuffled-based resampling procedure to assess the statistical significance of the SC score to avoid false cluster discovery. The test relies on creating 1000’s of surrogate datasets through a shuffling or randomization procedure that is applied to the row/columns of the single neuron SSIM matrices. This procedure effectively destroys any dependencies among the neuron’s single neuron SSIM matrices and, consequently, the associated clusters with the CS neuron map (Fig. 4B). The empirical SC score is only considered statistically meaningful if it falls outside a distribution of surrogate SC values calculated from the 1000’s of surrogate datasets created in this manner (Fig. 4C, see blue dots; for further insights on conditional resampling see (Amarasingham et al., 2012; Schemper, 1984). This unsupervised cluster detection and validation procedure supports the identification of statistically meaningful clusters of computationally similar neurons within the CS neuron maps.

## 3.0 Results

We apply the SIMNETS analysis framework to four different datasets. First, we apply the algorithm to a population of simulated neurons for which spike patterns were constructed to simulate three subnetworks encoding information using firing rates, precise spike timing, or a combination of the two (Fig. 5). Next, we also apply it to three publicly available *in-vivo* experimental population recordings datasets from (1) macaque primary visual cortex (Kohn and Smith, 2016; Teeters and Sommer, 2009), (2) macaque primary motor cortex (Rao and Donoghue, 2014), and (3) rat CA1 hippocampal region field (Pastalkova et al., 2015; Wang et al., 2015) to illustrate the types of results the SIMNETS framework produces and how they may be leveraged for further exploratory analyses and hypothesis testing (Fig. 6 – 11; for algorithm inputs/outputs for each dataset see: Table S1). Finally, we highlight some of the positive attributes of the SIMNETS analysis framework and Toolbox that are particularly advantageous when analyzing large-scale neural recording data (Fig. 12).

**Fig. 5.**
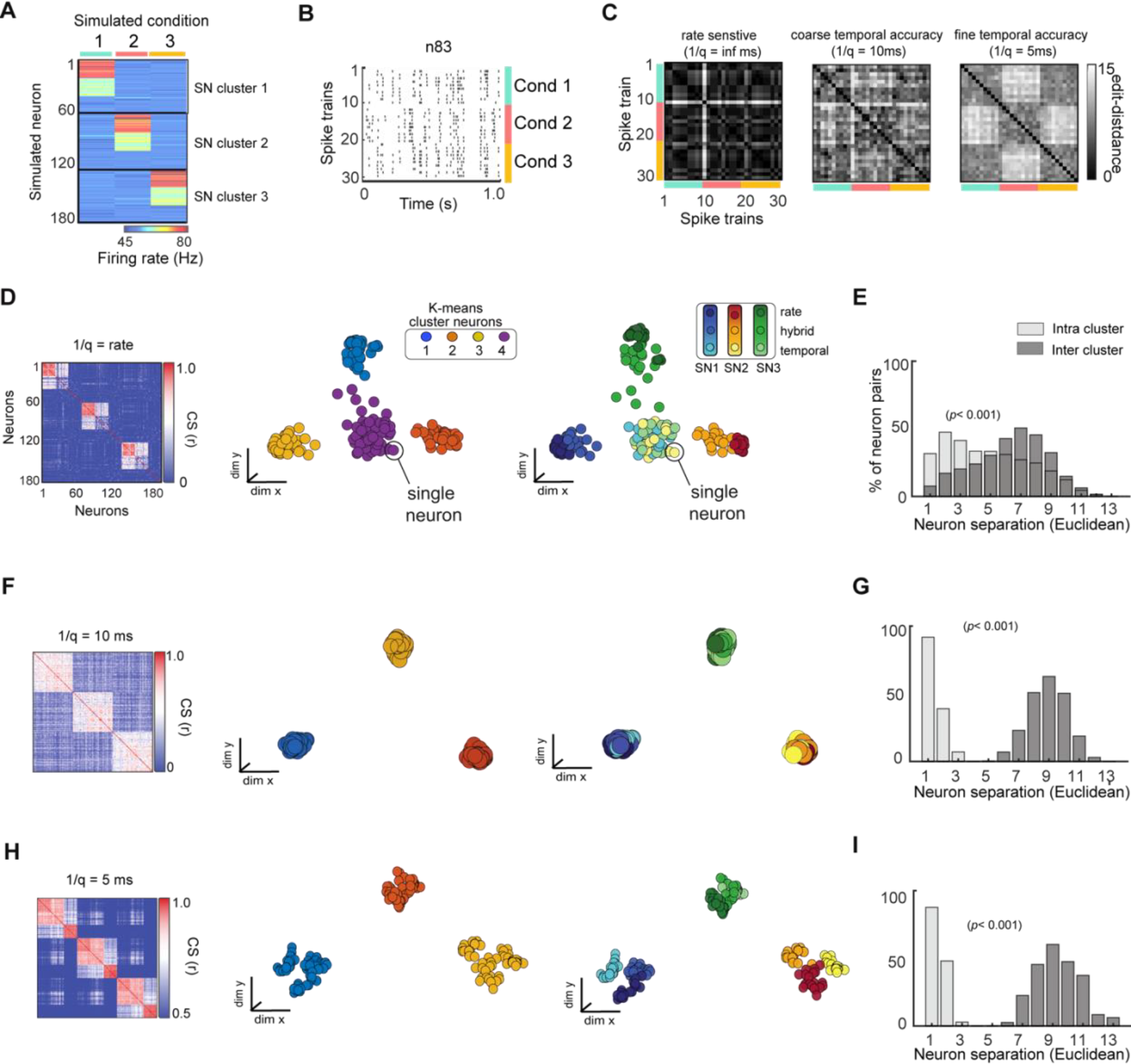
SIMNETS analysis framework successfully organizes a population of simulated neurons utilizing heterogeneous spike train encoding schemes into their ground-truth CS neuron clustering. **A)** Population representation of spike trains from a population of 180 simulated neurons with firing rate constructed according to three different coding schemes (rate, temporal, mixed); 10 spike trains for each of three conditions (C1-3) for an ensemble of N = 180 simulated neurons. The neurons show a distinct pattern of spike rates across conditions (red, green, or blue, color bar), illustrating the different subnetwork groups. **B)** Raster plot for one example neuron (n83) to illustrate a hybrid encoding scheme in which there is both rate change and different temporal structure in response to each of the conditions (C1-blue, C2-red, C3-yellow). **C)** Three SSIM matrix representations for neuron #83, illustrating trial similarities organized according to stimulus blocks, illustrating the effect of q that emphasized firing rate (left; (1/q): infinite), coarse temporal (middle; 100 ms (coarse), and 5ms (fine) temporal differences in spike pattern on each train. Note that the coarse and fine temporal matrices best reveal this neuron’s condition-dependent activity patterns. (**D**, **F**, **H**) Population CS matrices for the population to identify subnetworks using pure rate codes (q = 0, no cost for shifting spikes in time), coarse temporal accuracy (1/q = 100 ms), or fine temporal accuracy (1/q = 5ms). Neurons are colored according to clusters identified using k-means (left) or ground truth subnetwork designation (right). **E**, **G**, **I)** Histograms showing normalized separation between neurons within each of the different SIMNETS maps for ground-truth “Intra-cluster” neuron pairs (light gray) and “Inter-cluster” pairs (dark grey).

**Fig. 6.**
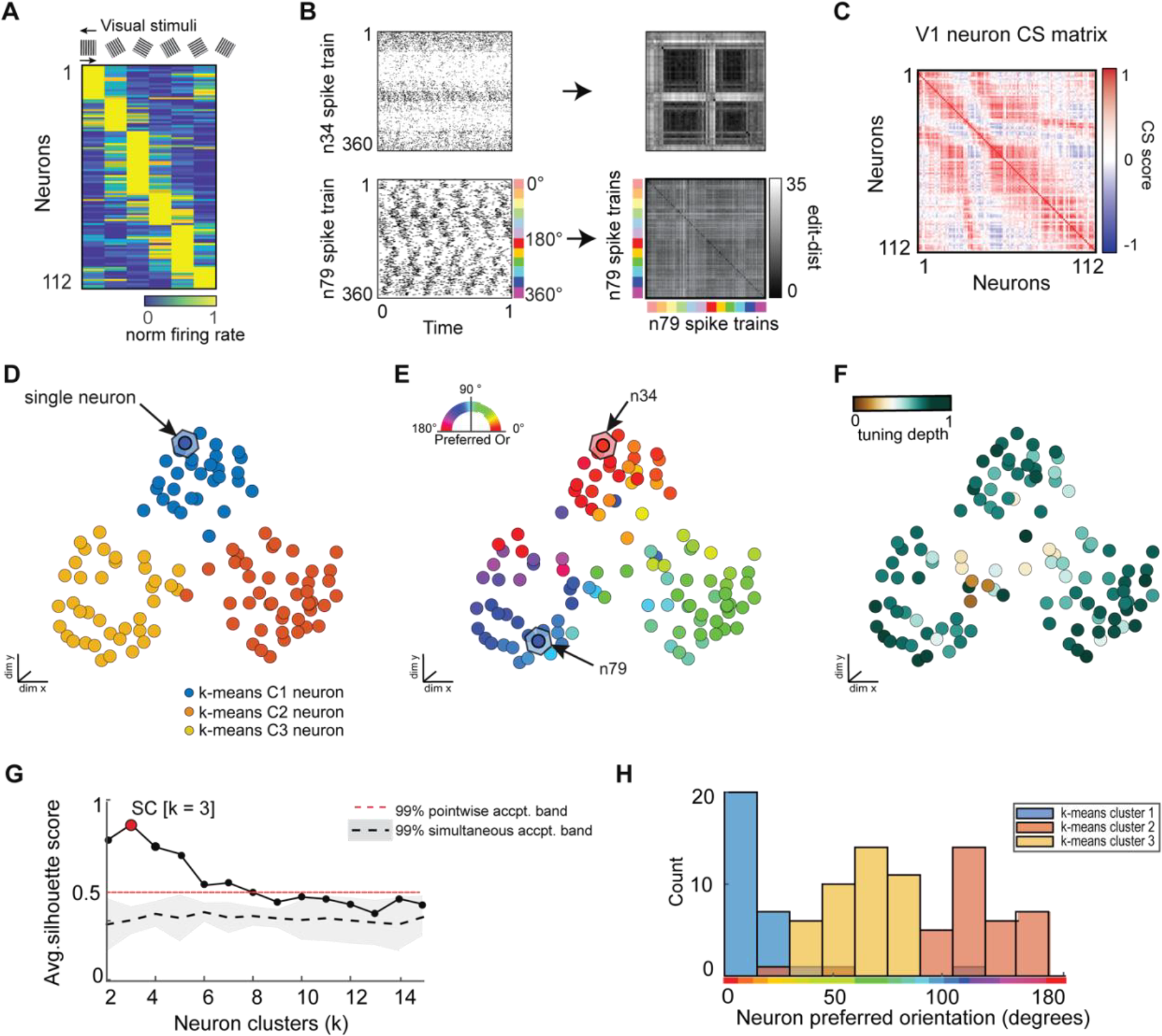
SIMNETS analysis framework captures known computational properties among a population of primate visual cortex neurons without the need for extrinsic labels or tuning models. **A**) Normalized trial-averaged firing rates of a population of V1 neurons (N = 112, neurons) during the presentation of 12 different drifting grating stimuli (S = 360 spike trains) for 1.28 s at six different orientations: 0, 60, 90, 120, 150 degrees, and two drift directions (rightward and leftward drift). Neurons are ordered along the y-axis according to peak firing rate for visualization purposes, only. **B**) Two example single neuron raster plots (n34 and n79) and their corresponding SxS single neuron SSIM matrices for VP (q = 35). The colored line indicates the stimulus orientation and drift direction. **C**) SIMNETS NxN CS map. **D**-**F**) Low-dimensional (3xd) population CS map, with neurons labeled according to k-means cluster assignments (**D**), the neuron’s preferred stimulus orientation (**E**), and their orientation tuning strength (**F**). Example neurons, n34 and n79 (**B**), are indicated with arrows in (**E**). **G**) Average silhouette score for the population CS map as a function of the number of clusters. Red circle indicates the SC at k = 3 (SC = 0.89; p<0.0001; shuffled data). H) Histogram showing the preferred orientation of the neurons within each of the three k-means CS neuron clusters from (**D**).

### Simulated Neuron Population

We evaluated the performance of the SIMNETS algorithm using simulated neural data with known ground truth. This allowed us to verify that the algorithm clusters neurons with similar informational content even when they use different encoding schemes. We simulated a population of 180 neurons (N) composed of three computationally distinct subnetworks, SN1, SN2, and SN3 (Fig. 5A). We generated an output structure for these subnetworks across three simulated experimental conditions (C1, C2, and C3). Each subnetwork emitted a similar “baseline pattern” for two of the conditions and a different pattern for a unique “preferred” condition (C1 for SN1, C2 for SN2, and C3 for SN3). Neurons from a common subnetwork were designed to encode similar information (i.e., respond differentially to one specific condition) using different encoding formats; for example, 20 neurons in SN1 encoded condition A through a change in spike rates (rate-coding), another set of 20 neurons in the same subnetwork responded with a change in the timing of their spikes (temporal-coding neurons), while the remaining neurons responded with changes in both spike rates and the spike time variations (mixed coding) (e.g. Fig. 5B). The other two subnetworks were also generated to include examples of rate coding, temporal coding, and mixed coding artificial neurons in the same proportions. We simulated 30, one-second-long spike trains for each of the 180 artificial neurons, which included ten repetitions of each condition (S = 30 spike trains per neuron) (See Table S1 for analysis summary).

The SIMNETS algorithm was applied using three different temporal accuracy values for the VP spike train similarity metric: a pure rate code (q = 0, i.e., no added cost for shifting spikes in time), 100ms (q = 10), 5ms (q = 200). The q parameter determines how sensitive the similarity metric is to differences in the timing of individual spikes, by setting an upper limit on how far spikes can be shifted in time (See Fig. 5C, for three different single neuron SSIM matrix). With a setting of q = 0, the neurons operating with a rate-based encoding scheme and mixed coding neurons are grouped into three functionally distinct clusters in the CS map (Fig. 5D, 2^nd^ column), while temporal coding neurons form a single cluster at the center of the map (Fig. 5D, 3^rd^ column). As the value of q increases, the VP algorithm “cost” function becomes sensitive to differences in spike timing in addition to the total number of spikes (Victor and Purpura, 1996). SIMNETS correctly groups all neurons into three distinct clusters that reflect the ground-truth functional ensemble assignments (Fig. 5F). At the highest q value (Fig. 5H), the CS map shows sub-groupings within each of the three ground-truth subnetworks that correspond to neuron’s encoding formats (Fig. 5H, 3^rd^ column); however, the optimal number of clusters remains in agreement with the ground-truth functional subnetwork assignments (K_sc_ = 3). By specifying a higher partition value for the k-means clustering step of the algorithm (e.g., k = 9), the sub-groupings within the detected clusters are defined by the coding properties of the neurons (data not shown). For a demonstration of the interaction between the cluster number and the SIMNETS hyperparameters, perplexity, and q, see Supplementary Fig. S2.

We compared the distribution of interneuron separation (Euclidean distance between neurons in the SIMNETS in the CS map) for neuron pairs within and between the artificially generated (ground truth) subnetworks (Fig. 5 E, G, I, histogram). The within-subnetwork similarity was significantly higher than between-subnetwork values in all cases (Mann-Whitney p<0.001). For q values > 0, there was no overlap between the two distributions, indicating the complete separation of the artificially generated subnetworks. Our results demonstrate that the SIMNETS algorithm can accurately separate neurons according to their computational properties, even if they employ different coding schemes to represent information using spike rates or temporal patterns.

### Application to Empirical Multi-neuronal Recordings

We next applied SIMNETS to three distinct datasets of simultaneously-recorded, multiple single-unit recordings from (1) nonhuman primate (NHP) primary visual cortex (V1) during visual stimulation (Kohn and Smith, 2016; Smith and Kohn, 2008), (2) NHP motor cortex (M1) neural recordings during a center-out reaching task (Rao and Donoghue, 2014), and (3) rat CA1 hippocampal region recordings during a left-right alternation maze task (Pastalkova et al., 2015, 2008). These datasets were chosen because the functional roles of the neurons in these areas have been reasonably well-established and the information content of the single neuron and ensemble spiking patterns extensively described (Gur and Snodderly, 2007; Livingstone and Hubel, 1984; Rao and Donoghue, 2014; Smith and Kohn, 2008; Truccolo et al., 2008).

We note that these results are meant to illustrate the capabilities and utilities of the SIMNETS analysis pipeline and are not intended to represent a full characterization of the role of any identified putative subnetworks. For a full analysis and interpretation, many more datasets (sessions/subjects) would be required, as would various parameter sweeps to evaluate computational processes.

### Macaque V1 Neural Population Recordings during Visual Stimulation

We analyzed a dataset of 112 simultaneously recorded V1 neurons (recorded using a 96- channel “Utah” electrode array) during the presentation of drifting sinusoidal gratings in an anesthetized Macaque monkey (Kohn and Smith, 2016) (Fig. 6A). We extracted 1 second of spiking data from the first 30 repetitions of each stimulus (S = 360), starting 0.28 seconds after stimulus onset (see Fig. 6B for raster plots and single neuron SSIM matrices for two example neurons) to capture the neural stimulus response period (eliminating response delay). SIMNETS was applied to these NxS spike trains generated with a fixed temporal accuracy setting of 1/*q* = 0 ms. The resulting NxN CS neuron matrix is projected into a lower-dimensional CS space upon which further statistical and clustering analyses are performed (Fig. 6D - F). An unsupervised k-means cluster analysis identified 3 clusters of related single neurons in the V1 CS map (Fig 6D, G; SC = 0.78, peak max average silhouette value), indicating that computational features did not fall along a functional continuum but instead were organized into three functionally distinct putative subnetworks.

To validate the performance of the SIMNETS algorithm, we examined the relationship between the structure of the CS map with respect to the known tuning properties of V1 neurons. We used classic parametric tuning models to estimate the receptive field properties of all neurons and labeled the neurons in the CS neuron map according to their “preferred” stimulus orientation (Fig. 6E) and tuning depth (Fig. 6F). Each neuron’s preferred orientation was calculated by fitting a gaussian distribution to the stimulus-dependent firing rates and finding the orientation that maximizes the function over the range of orientations angles (θ = [0,180)) across both left and right drift directions while tuning strength was calculated as the normalized difference between the peak and trough of the tuning function (See Supplementary Methods). Again, we emphasize that the SIMNETS procedure is an entirely unsupervised approach and does not rely on any knowledge of the experimentally induced neuron tuning functions to generate the computational similarity map or to identify putative CS neuron clusters therein. We simply labeled each neuron with values obtained from its relationship to an experimental variable (here, grating direction) after the neurons were plotted in SIMNETS computational space. A pairwise circular-linear correlation (*r*_cl_) statistical analysis revealed a strong positive correlation between neurons’ separation in the low-dimensional CS neuron space and the differences in their preferred orientation parameters (Pearson, *r*_cl_ = 0.89), which confirmed that neurons with similar preferred orientation were generally mapped near one another (Fig 6E). The exception to this trend was observed for neurons with weaker tuning strengths (Fig 6F, brown points), which tended to aggregate at the center of the CS neuron map. Overall, the distribution of the neurons’ preferred orientations corresponded well with the three observed clusters in the CS neuron space (Fig 6H). Collectively, these results confirm that the neurons’ sensitivity to the different stimulus orientations is a major contributor to the structure of the V1 computational architecture and reveals the presence of three computationally distinct groups of neurons within the context of this experimental paradigm.

The overall agreement between known computational features in V1 and the computational map generated using SIMNETS suggests that the algorithm will be able to describe computational architectures for neural systems where the tuning function geometries are not known *a priori*. However, as previously mentioned, univariate parametric tuning models of V1 neuron function may only provide a limited picture of the neuron’s computational role within cortical networks; indeed, there are likely several additional shared computational features that might have contributed to the structure of the V1 CS neuron space – such as anatomical connectivity, receptive field location, spatial frequency selectivity, direction-of-motion selectivity, response reliability, and anesthesia-induced fluctuations of local excitatory-inhibitory synaptic inputs, etc. To better illustrate the relationships between the different computational geometry representations captured within the local structural features of the V1 CS neuron map, we used a multi-scale state-space analysis approach to visualize the computational geometry representations across different scales of organization within the CS network (Fig 7).

**Fig. 7.**
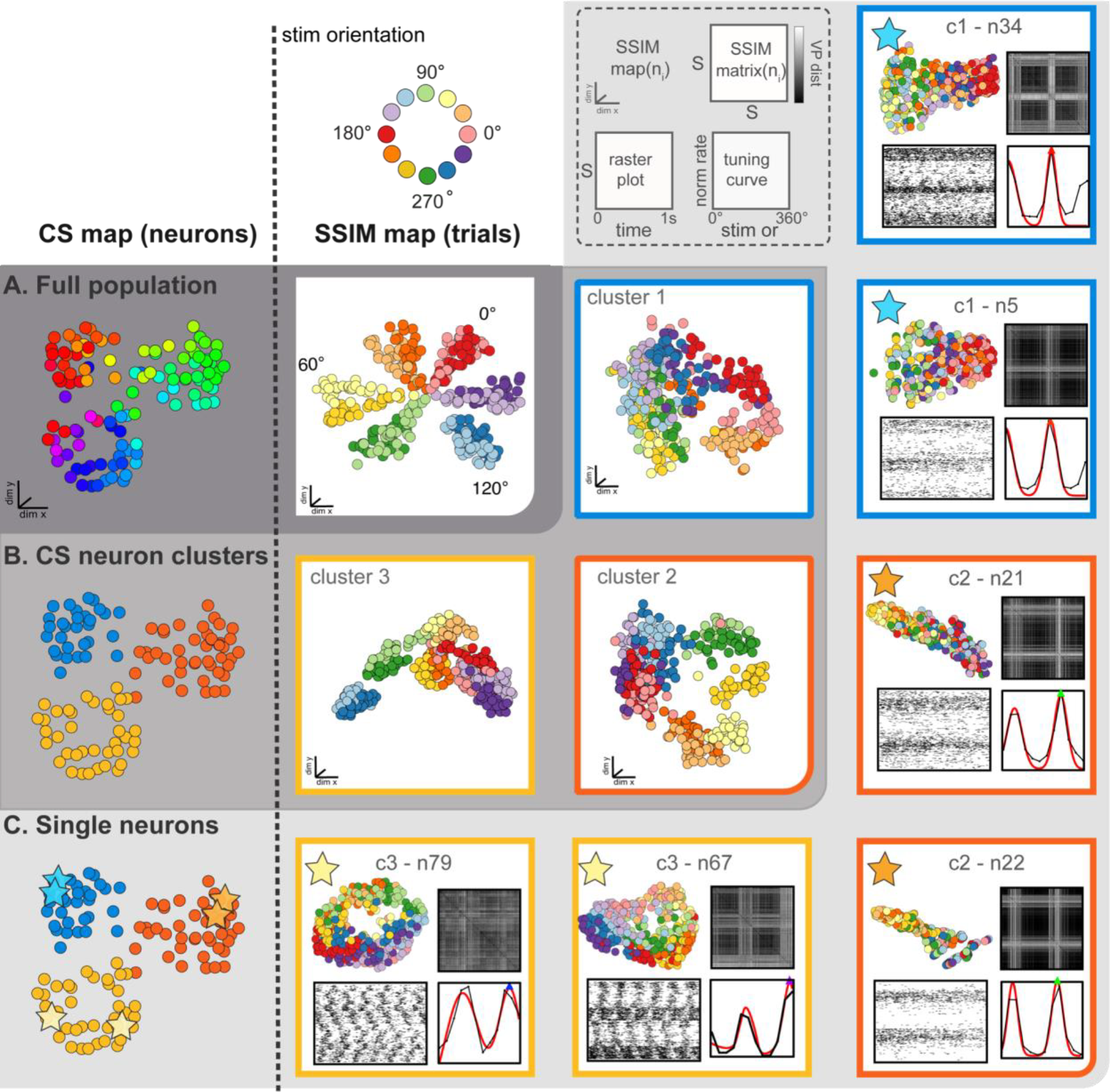
The SIMNETS framework facilitates the comparison of the computational geometry representations distributed within and across different levels of organization within the primate visual cortex dataset. **A)** hierarchical neural geometry distribution diagram for the V1 SIMNETS CS neuron map ((**A)**, left; dark gray) highlights the relationship between the population-level V1 neural computational geometry ((**A**), right) and that at lower levels of organization (CS neuron clusters; (**B**)) and single neuron examples (**C**)). All maps shown to the left of the broken vertical line are CS maps (each point represents a neuron); maps shown to the right of the broken line are multi-level SSIM maps (points are trials). **A**) population-level V1 CS neuron map with neurons labeled according to preferred orientation (left; same as Fig. 6) and corresponding population-level SSIM map (right) shows the neural computational geometry for the full V1 population. Each point in the SSIM map corresponds to a single trial; pairwise distances between points correspond to the dissimilarity among the V1 population spiking pattern on those trials. Colors correspond to the drifting grating orientation (Or); light vs. dark hues correspond to right (Or <180°) or left drift direction (Or >180°). **B)** Left: SIMNETS CS neuron map with neurons labeled according to their assigned CS neuron clusters (points are neurons). Right: the corresponding mesoscale CS cluster SSIMS representations (right; points are trials) for each of the three identified k-means clusters (blue, cluster 1; orange, cluster 2; yellow, cluster 3). **C)** Left: SIMNETS CS neuron map indicating the location of the example neurons (stars). Right: a single neuron SSIMS map, matrix, spike train raster plot, and parametric orientation tuning function are shown for two example neurons from each of the three CS neuron clusters.

The DR techniques used to visualize the computational geometry representations of the single neuron SSIM maps (introduced in Fig. 2) can be expanded to represent the combined output space of multiple neurons simultaneously, such as the CS neuron clusters (i.e., putative subnetworks) or the neuron population. For multi-neuron SSIM maps, each point corresponds to the collective spiking pattern generated by all neurons within an ensemble (e.g., CS neuron cluster or full population) on a single trial, and the distance between points corresponds to the difference (spike train distance) in the neurons collective spike pattern across time. Here, we use this approach to visualize the computational geometry structure for the full neuron ensemble (Fig. 7A), the three CS neuron clusters (Fig. 7B), and two example single neurons from each of the three CS neuron clusters (Fig. 7C). Standard single neuron plots (parametric tuning functions and rasters) were also included for comparison.

### Macaque M1 Neural Population Recordings Center-out Reaching Task

As a second example, we applied the SIMNETS algorithm to a dataset of 103 M1 neurons recorded using a 96-channel electrode array in a macaque performing a planar, 8- direction arm reaching task (see Methods for more details). We extracted 1-second long spike trains (S = 114) from each neuron during all trials where the monkey successfully reached the cued target, starting 0.1 seconds before movement onset (Fig 8A; see Table S1 for algorithm inputs and parameters). The SIMNETS analysis revealed that the neurons were organized into 3 neuron clusters in the M1 CS map, with a peak silhouette value that was slightly less than that observed for the V1 dataset (Fig. 8D and G, SC score = 0.71, M1 peak average silhouette).

**Fig. 8.**
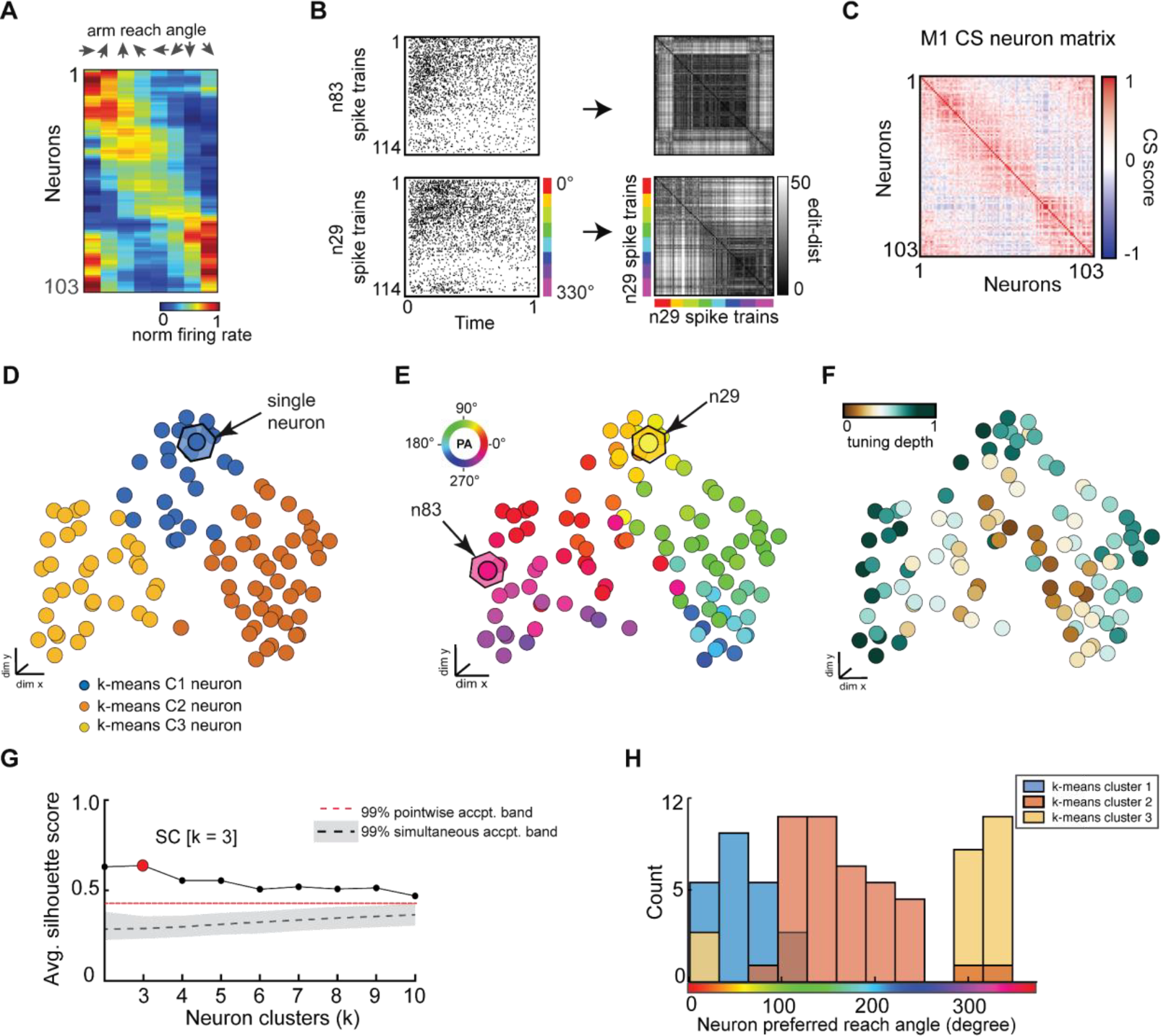
SIMNETS CS neuron map captures known computational properties among a population of primate motor cortex neurons without the need for extrinsic labels or tuning models. **A)** normalized trial-averaged firing rates for a population of simultaneously recorded M1 neurons (N = 103, neurons) as a function of reach direction for a planar, 8-directional reaching task. For visualization purposes, neurons were ordered along the y-axis according to their peak condition-dependent firing rates. Neurons are ordered along the y-axis according to peak firing rate for visualization purposes, only. **B**) Two example single neuron raster plots (n29 and n83) and their corresponding single neuron SSIM matrices for a VP edit-distance setting of q = 15 (i.e., 66 ms temporal precision). Colored line indicates the reach angle associated with each spike train. **C)** SIMNETS NxN CS matrix. **D - F**) Low-dimensional population CS map, with neurons labeled according to k-means cluster assignments (**D)**, the neuron’s preferred reach angle (**E**), and tuning strength (**F**). Example neurons, n29 and n83 (**B**), are indicated with arrows in (**E**). **G**) Average silhouette score for the population CS map as a function of the number of clusters. Red circle indicates the SC at k = 3 (SC = 0.64; p<0.0001; shuffled data). **H**) Histogram showing the preferred reach angle of all neurons within each of the three k-means CS neuron clusters (shown in **D**).

To examine the properties of the SIMNETS CS map, we characterized the neurons preferred reach angle (Fig 8F) and tuning depth (Fig 8F), in a similar manner to the V1 dataset. Each neuron’s preferred reach direction was estimated by fitting a von Mises distribution to the firing rates as a function of direction (Fig. S3) (Mardia and Zemroch, 1975) A circular-linear correlation (rcl) analysis between the neuron’s preferred reach direction and their inter-neuron separation in the CS map revealed a significant positive relationship (Pearson, rcl = 0.92; p = 0.001), confirming that neurons with similar preferred directions were clustered within near-by regions of the map and neurons with dissimilar preferred directions were spatially distant (Fig 8D-E, right plot). After plotting the distribution of preferred reach direction, we again observed that the neuron tuning properties were similar across each of the three detected CS neuron clusters (Fig 8H).

SIMNETS analysis suggests that the computational properties of M1 neurons are not uniformly distributed along a functional continuum, as evidenced by the distinct CS neuron clusters identified in the population CS space. The separation between the CS neuron clusters is most apparent for CS neuron cluster 2 and cluster 3 (Fig. 8D, orange and yellow CS clusters), which mirrors the discontinuity in the distribution of preferred reach angles (Fig. 8E). These results agree with previous findings, which support the hypothesis that the biomechanical constraints of the limb are reflected in an uneven distribution of preferred directions among motor cortical neurons (Lillicrap and Scott, 2013). The identified CS neuron clusters may reflect functional modules for other coding features such as joint angles, which are not evident in traditional rate-based, raster histogram analysis or population coding methods.

As with the V1 dataset, we used a multi-scale latent space analysis to visualize the computational geometry across different scales within the CS network from the full ensemble (Fig. 9A) to CS neuron clusters indicative of potential subnetworks (Fig. 9B) and individual neurons functional properties (Fig. 9C). Standard single neuron plots (parametric tuning functions and raster plots) are again included for comparison (Fig. 9C). Each M1 neuron had a unique single neuron SSIM map that captured several computationally relevant features of the neurons’ spike train outputs (Fig 9C; SSIM matrices/maps for two example neurons). In contrast to the regular conical and circular parabolic shapes of the V1 single neuron SSIM spaces (Fig 7C), the M1 SSIM maps displayed more irregular and heterogenous global geometries (Fig 9C).

**Fig. 9.**
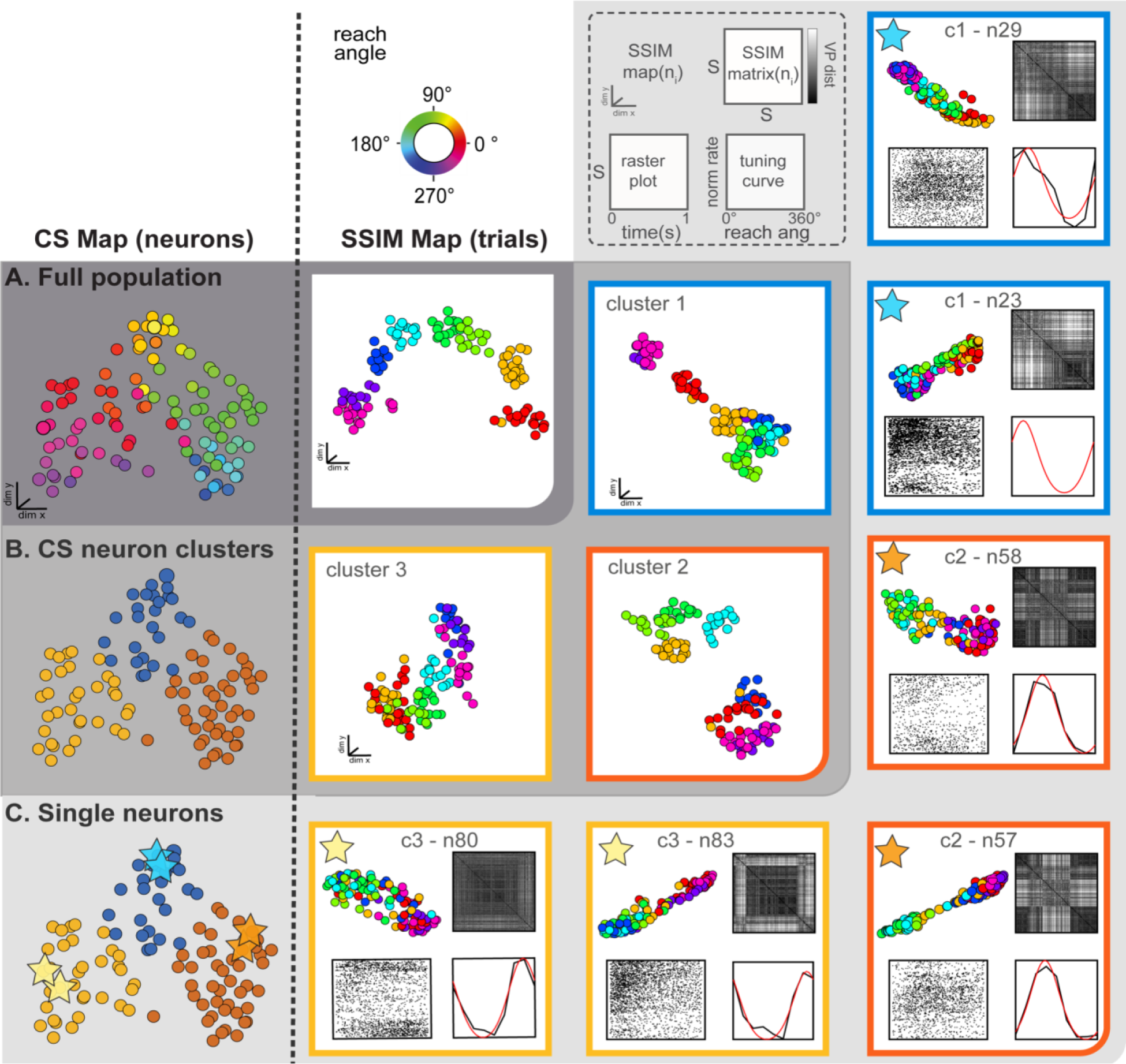
The SIMNETS framework facilitates a comparison of the computational geometry representations distributed across multiple scales of organization within the primate motor cortex test dataset. Illustrative guide on how M1 SIMNETS CS network map (left of broken black line) may be used to exploring the neural geometry representations (right of broken black line) at the population level (**A**), the CS cluster level (**B**) and single neuron level (**C**) (layout same as for Fig. 7). **A)** Left: SIMNETS M1 CS neuron map for full neuron population, with neurons tuning property labels (preferred reach angle). Right: SSIM maps for full population, with trial condition labels (reach angle). **B**) Left: SIMNETS M1 neuron map with CS neuron cluster labels. Right: separate cluster SSIMS neural subspace maps for each of the CS neuron clusters. **C)** Coding features of two example single neurons from each of the three CS neuron clusters (indicated by stars in SIMNETS CS neuron map). Each single cell panel shows a single neuron SSIMS map for one cell (top left) and a single neuron SSIMS matrix (top right); a raster plot for the same data for that cell (bottom left); reach angle tuning function for that cell (bottom right). Single neuron SSIM maps/matrices are similar within CS neuron clusters (**C**) and appear to mirror their corresponding CS cluster neural geometries at the mesoscale-level (**B**).

### SIMNETS Subnetworks in a Population of Rat Hippocampal CA1 Neurons

Lastly, we used SIMNETS to reveal the computational organization of a population of hippocampal neurons, a region that is both functionally and structurally quite different from the primary visual and motor neocortical areas examined above. We applied SIMNETS to a publicly available dataset of rat CA1 hippocampal neurons recorded using multi-site silicon probes in a rat performing a left/right-alternation navigation task in a “*figure*-8” maze (Pastalkova et al., 2015) (Fig. 10A). All neurons with firing rates greater than 5Hz were included in the analysis (N = 80/106 recorded neurons). The rat performed 17 correct trials (T = 17, trials), taking an average of 4.3 seconds to reach the reward location at the end of the left or right arm of the maze. Spike trains used for the SIMNETS algorithm were obtained by extracting 1-second of spiking activity for each 80cm increment of distance traveled by the rat (Fig 10B, *left*, example single neuron raster plots with S spike trains; see Supplementary Methods in Supp. Info. document). This resulted in a set of S = 442 spike trains (17 trials x 26 time-epochs) per neuron.

**Fig. 10.**
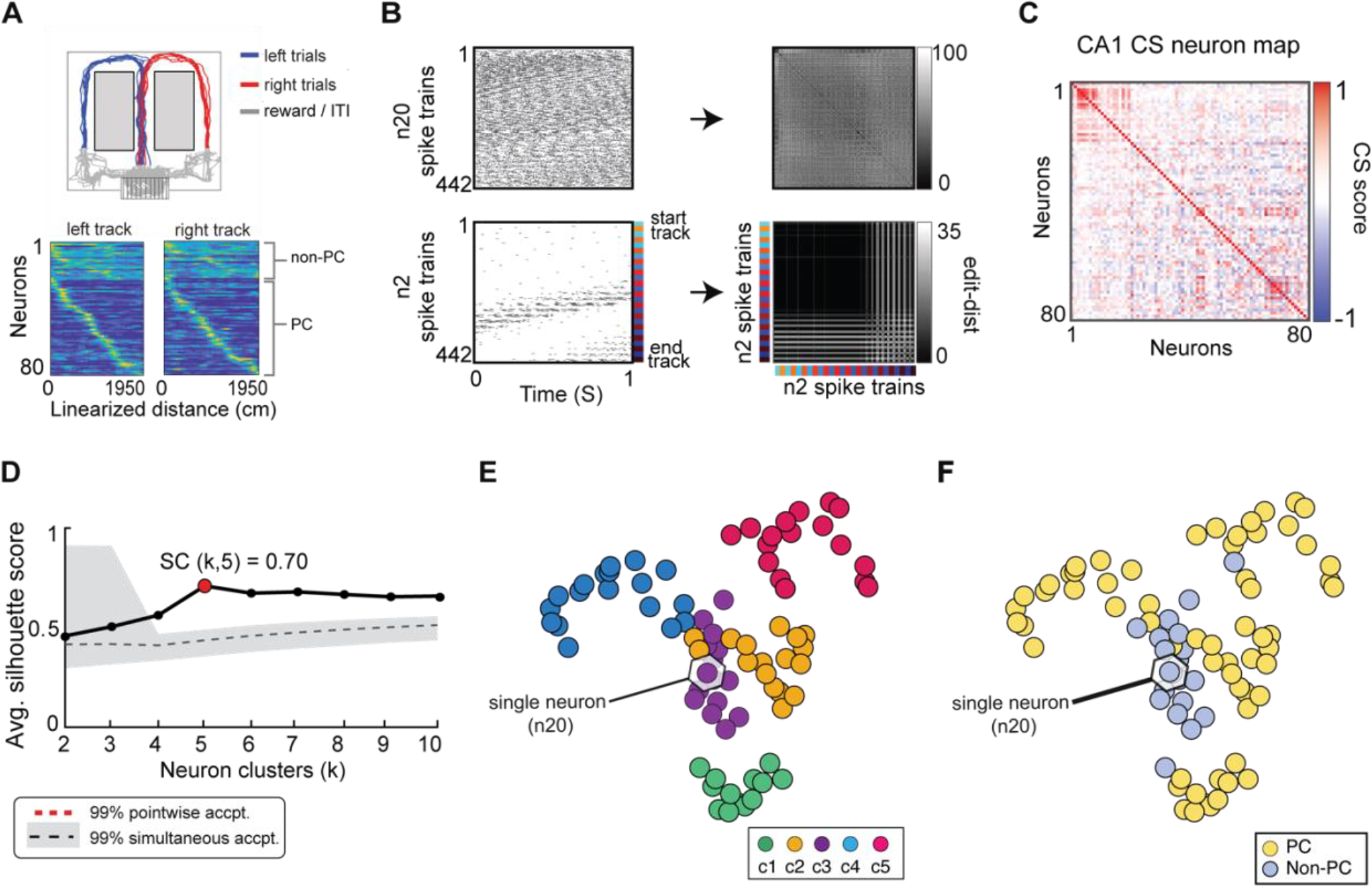
SIMNETS analysis framework captures the known computational relationships among a population of Hippocampal CA1 neurons in an unsupervised manner. **A)** Task description: “figure-8” maze showing rat’s position during left-right alternation task (top). Red and blue lines show the rat’s location during correct right and left trials (T = 17, trials), respectively. Gray lines show the rat’s location during reward and inter-trial interval periods. below: normalized firing rates are shown for each neuron (N = 80, neurons) as a function of linearized distance on track (50 cm bins) during left and right trials (below). For visualization purposes, neurons were ordered according to the latency of their peak response along the track and according to their characterization as a non-place cell (non-P.C, N = 20, light blue) or a place cell (P.C.; N = 60, yellow). **B)** Two example single neuron raster plots (n7 and n20) showing S=103 spike trains of 1-second duration (26 spike train segments per T trials) and their corresponding SxS single neuron SSIM matrices for VP [q = 35]. Colored line/dots indicate the rat’s location. **C**) SIMNETS NxN CS matrix. **D)** Average silhouette score for the population CS map as a function of the number of clusters. Red circle indicates the SC at k = 5 (SC = 70; p<0.0001; shuffled data). **E-F)** Low-dimensional (3xd) population CS map, with neurons labeled according to Ksc = 5 k-means cluster assignments (**E**), and the neuron’s place-cell like firing properties (**F**) (P.C., blue dots; non-P.C., grey dots).

SIMNETS was applied to the set of SxN neurons, using the VP temporal sensitivity setting of q = 35 and perplexity of perp = 10 (Fig. 10B - C; see Table S1 for all algorithm inputs/outputs). After generating the low-dimensional CA1 population CS neuron map, we characterized the organizing properties of the neuronal relationships using the unsupervised k-mean cluster detection/validation procedure. This procedure identified Ksc = 5 statistically significant CS neuron clusters within the CA1 CS neuron map (SC = 0.70, peak max average silhouette value; Fig. 10D), suggesting that the neurons are organized into five computationally distinct clusters of neurons. To verify this, we used two different classic functional characterization approaches to assign functional ids to the neurons within the SIMNETS CS map and assess their functional organization within the context of the CS neuron map (Fig, 10F). Specifically, we categorize the CA1 neurons as having place cell-like activity (n = 60, place-cells, PCs) or lacking place cell-like activity (n = 20, non-place-cells, non-PCs) using an information-theoretic measure (Skaggs et al., 1992) and a spatial firing rate tuning functions (Fig. 10B, example single neuron raster plots; see Supplementary Methods for analysis details). As is reported in other studies, some of the neurons categorized as place-cells exhibited complex firing responses, such as multiple peak place-dependent firing responses (Fig. S3D, show example single neuron response functions). An examination of the physiological properties of the identified k-means neuron clusters revealed that one cluster was almost entirely (95%) composed of non-PCs (Fig 10E-F, k-means cluster c3, purple), while the other four k-means CS neuron clusters were entirely or almost entirely (>92%) composed of PC’s (Fig 10E-F). This indicated one of at least three possibilities: 1) the “non-PC CS cluster”, cluster c3 corresponded to a cluster of untuned, “uninformative” neurons, 2) that the individual neurons computed a common task-related signal that was unrelated to place-coding, or 3) that place-coding signals were only apparent at the level of the neuronal cluster rather than the individual neurons (Fig. 10 E, cluster c3). While a full exploration of these different possibilities is beyond the scope of this work, we do provide a demonstration of how the SIMNETS CS neuron maps may be used to visualize the neural computational geometry across scales for hypothesis formation and testing (Fig. 11).

**Fig. 11.**
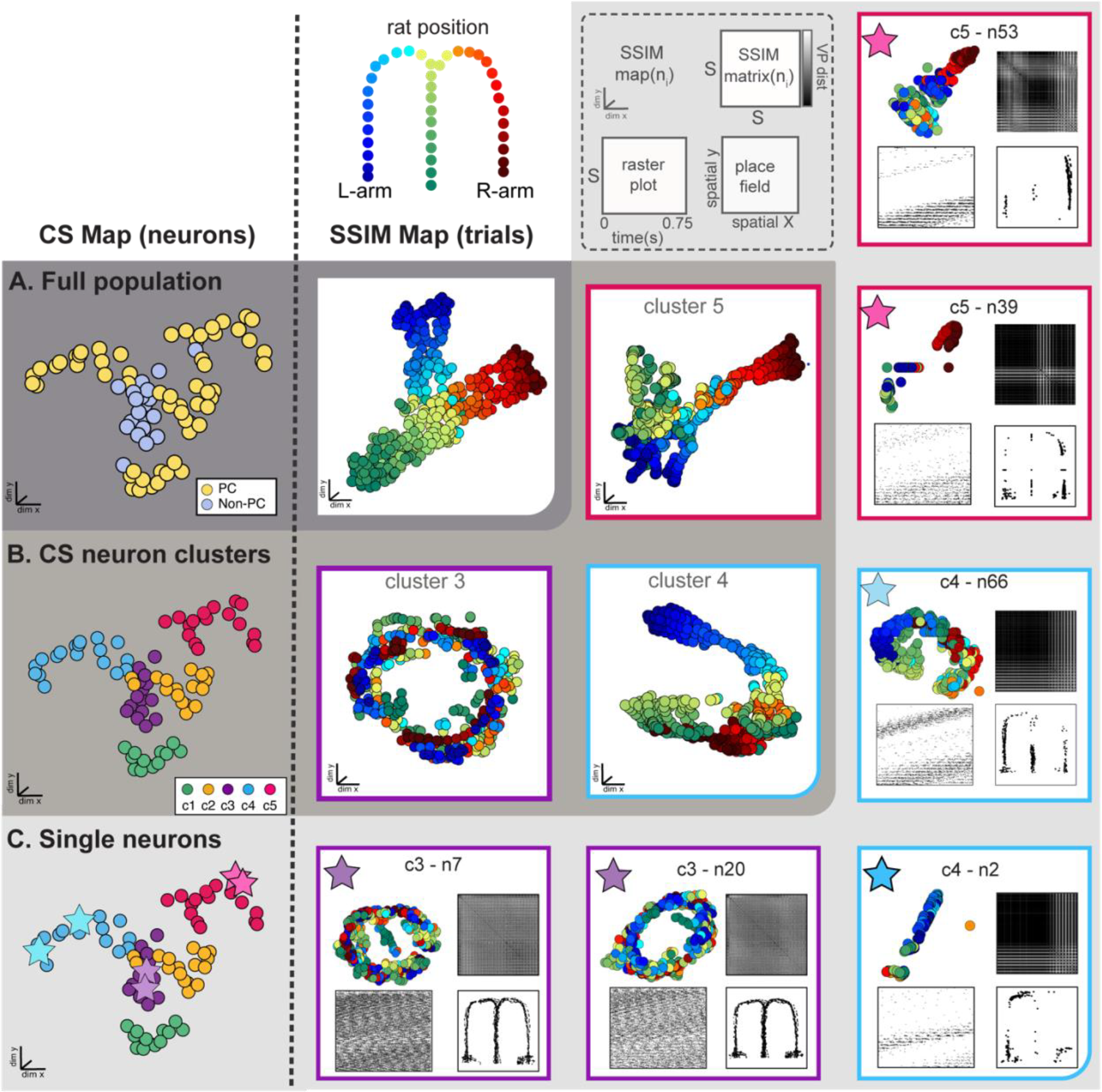
SIMNETS framework facilitates a comparison of the computational geometry representations distributed across multiple scales of organization within the hippocampal CA1 test population. SIMNETS map (left of broken line) and SSIM map (right of broken line) are shown for the CA1 neuron population recording during left-right alternation behavioral task. **A**) Left: SIMNETS population CS neuron map for rat CA1 neuron population recording during left-right alternation behavioral task. Yellow points correspond to neurons with place-cell activity (PC). Right of broken line: population SSIM neural subspace map showing single trial CA1 neuron population spike patterns. Colored points represent the rat’s spatial location in the maze. **B**) Left: SIMNETS CS neuron map with CS neuron cluster labels, with neurons as points. Right: three separate cluster-level SSIMS neural subspace maps (points correspond to spike trains) for CS neuron clusters c3, c4, and c5 (see Fig. S3C for other clusters). **C)** Right of broken line, in light gray: different single neuron representations, two example single neurons from each of the three CS neuron clusters (indicated by stars in SIMNETS CS neuron map). Each single cell panel shows a single neuron SSIMS map for one cell (top left) and an SxS single neuron SSIMS matrix (top right); a raster plot for the same data for that cell (bottom left); single neuron spike field spatial map of maze (bottom right). Single neuron SSIMS maps/matrices are similar within CS neuron clusters and distinct across CS neuron clusters(**C**), and the different CS cluster SSIMS neural subspace maps (**B**) correspond to the spatial differences between CS neuron clusters in the SIMNETS CS map in ((**A**), left).

To better understand the computational properties of the neurons within and across each of the CS neuron clusters, we again examined the SIMNETS CS neuron map using spike train relational data to depict latent spaces representations different organizational scales, including the population-level (Fig. 11A, *right of broken line, dark gray panel*), mesoscale cluster-level (Fig. 11B, *right of broken line, medium gray panel*), and neuron-level (Fig. 11C, *right of broken line, light gray panels*). As expected, the spatial layout of the task-space was reflected in the topology of the population-level SSIMS map (Fig.11A, *right of broken line*, SSIMS map; see Fig. S3B, for different viewing angle), while each cluster-level SSIMS maps captured the place-dependent modulation of the clusters spiking patterns as the rat traversed a specific region of the maze (Fig. 11B, Fig. S3C). Surprisingly, the “non-PC neuron cluster”, (Fig. 11B, cluster c3) did not appear to reflect a cluster of non-responsive or “uninformative” neurons; instead, it had a toroidal shape that resulted from at least two dominant patterns of spike pattern modulation: a within-trial periodic spiking pattern in the x-y plane (Fig. 11B, cluster c3, x-y viewing angle) and a location or distance-dependent variance along the z-axis (See Fig. S3C, for x-z viewing angle). This torus topology was also observed in the single neuron SSIM maps of the neurons within cluster c3 (see Fig. 11C, purple box/stars, neurons c3-n7 and c3-n20), however, the distance-dependent x-z variance was less apparent at the single neuron level. Overall, this suggests that the activity of the non-PC neuron cluster, c3, displays dynamics that may reflect a different task variable that is in some way correlated with the rat’s performance on the task (e.g., speed profile along maze arms) or, alternatively, the intrinsic circuit dynamics (Pastalkova et al., 2008; Richard et al., 2013; Wang et al., 2015). Further investigation of this phenomenon is outside of the scope of the present work but is an example of how SIMNETS may be used to quickly uncover different types of functional relationships among neural representations that might not be otherwise evident when using classic population and single analyses. These results highlight the advantages of applying SIMNETS to neural recordings where tuning properties or other coded variables are not readily apparent or unknown *a priori*.

### Computational Efficiency and Analysis Run-time

The SIMNETS framework can rapidly analyze large numbers of neurons using a standard personal computer (e.g., 6-Core Intel Core i7, 64GB of RAM). With the current Matlab® implementation (see included SIMNETS software package), the simulated neuron population (N = 180, S = 30) and Macaque M1 neuron population (N = 103, S = 360) were processed in less than 2 seconds. Larger datasets, such as the Macaque V1 neuron population (N = 112; S = 360) and rat hippocampal CA1 neuron population (N = 80, S = 442), take approximately 10 seconds to process. This is significantly faster than a typical cross-correlation-based functional connectivity analysis. For example, a dataset of 1000 neurons can take an hour to processes with a standard cross-correlation analysis (Fig. 12A, blue line; utilzing Matlab® xcorr function), whereas out implementation of the SIMNETS algorithm takes only about a minute to process the same dataset (Fig. 12A, black line). The SIMNETS analysis is faster because it involves fewer spike train comparison operations (NxSxS) than the cross-correlations analysis (N2xSxS, across all time-lags), and the number of spike train comparisons per added neuron has a favorable linear scaling. If parallel or distributed computing resources are used to calculate the single neuron SSIM matrices, the execution time can be reduced even further to seconds or milliseconds. This would be especially useful for datasets with larger trial numbers (S), which are more computationally intensive than datasets with larger neuron numbers.

**Fig. 12.**
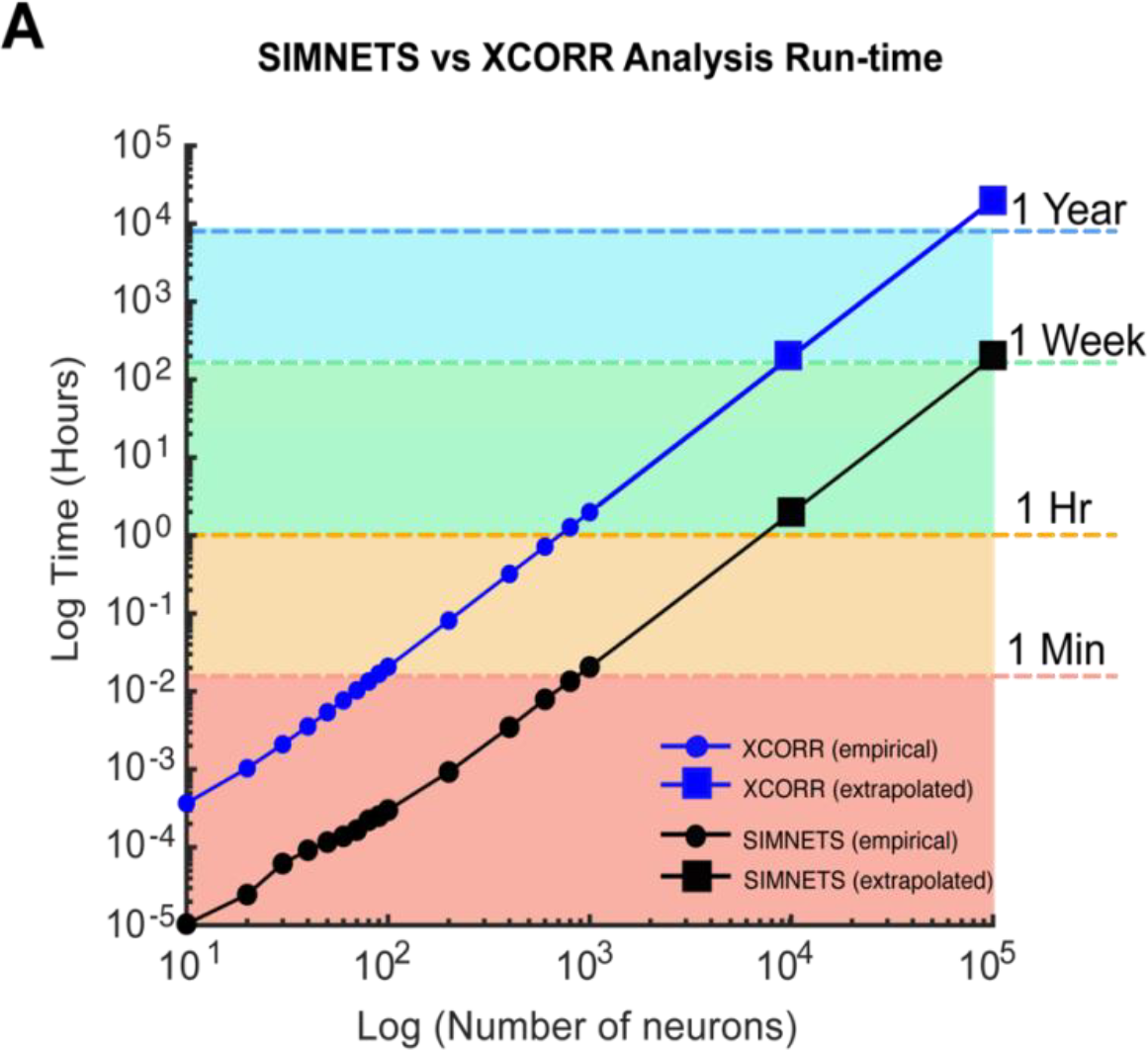
Computational run-time analysis for SIMNETS vs. cross-correlation algorithm. **A)** Log-Log plot showing SIMNETS algorithm (black lines) versus cross-correlation algorithm (XCORR, blue line) run-time as a function of neuron number (N = 10 - 100k, neurons). Empirical data points (round) are the mean run-time across 15 iterations. Extrapolated data points (square) were calculated from empirical data points for prohibitively large neuron population sizes (N = 10k - 100k). Colored regions indicate whether the population size was analyzed in under 1 minute (orange), 1 hour, 1 week, or 1 year. Software was run through MATLAB on a standard personal computer: 6-Core Intel Core i7 processor, with 64GB of RAM. Cross-correlation analysis relied on MATLAB’s xcorr function.

## 4.0 Discussion

Advances in multi-electrode recording technology have made it possible to record or image the activity of thousands of individual neurons simultaneously (Chung et al., 2022; Hsieh and Zheng, 2019; Paulk et al., 2022; Steinmetz et al., 2021; Vogt, 2019). The shift in focus from studying single neurons to population levels responses has brought about a new era of discovery, innovation, and collaboration (Buzsáki, 2004; Cunningham and Yu, 2014; Hurwitz et al., 2021; Kohn et al., 2020; Paninski and Cunningham, 2018; Paulk et al., 2022; Saxena and Cunningham, 2019; Urai et al., 2022); highlighting the need for new theoretical and analytical frameworks that can account for mesoscale computational processing operations and bridge the “representational gap” between neuron-level and population-level computational dynamics.

With this work, we introduced SIMNETS, a novel relational analysis framework designed to generate low-dimensional maps that provide a population-level view of computational similarity relations across sets of simultaneously recorded neurons. We tested and validated SIMNETS using a simulated neural population dataset. We then used three different experimental neural recording test datasets to demonstrate the general applicability of the analysis framework across a range of neural systems (visual, motor, cognitive), behaviors (awake or anesthetized), and different species (rat and macaque). This included a demonstration of a novel shuffle-based statistical procedure for identifying non-spurious CS neuron clusters within a low-dimensional population CS neuron map.

The critical difference SIMNETS and traditional functional connectivity measures is that we assess similarities among the neurons’ intrinsic latent space geometries, rather than the similarities of the neurons’ response time course. We emphasize that in contrast to neuronal functional connectivity analyses that measure time series correlations among the firing rates vectors or spiking patterns of *different* neurons (Abeles and Gat, 2001; Aertsen and Gerstein, 1985; Gerstein and Michalski, 1981; Grün et al., 2002b; Humphries, 2011; Kiani et al., 2015; Stuart et al., 2002). SIMNETS only directly compares spike trains generated by the same neuron, i.e., self-similarity. By quantifying the similarity of a neuron’s spike train outputs across all trials (a within-neuron 1^st^-order comparison), we generate a geometric representation of each neuron’s spike train output space, which we use as a “computational fingerprint”. The method recapitulated certain aspects of the neurons’ functional organization in an unsupervised manner, that is, without the use of trial labels or neuron functional labels (Kohn and Smith, 2016; Pastalkova et al., 2015; Rao and Donoghue, 2014).

SIMNETS projects the neuron population into a common coordinate space in a way that preserves the similarity relationships between the neuron’s computational fingerprints. The separation between neurons in the CS space provides a quantifiable measure of computational similarity, and CS neuron clusters reveal potential subnetworks with distinct computational properties. With the use of an appropriate dimensionality reduction method aims to preserves both local and global structure (e.g., PCA initialized t-SNE), the distances and spatial arrangement of the CS cluster can provide additional insight into the type of information that is being computed (Hinton and van der Maaten, 2008; Kobak and Linderman, 2021; Lee et al., 2015). SIMNETS could help resolve long debated issues as to whether classes of neurons that represent a unique set of features belong to a discrete computing module, or alternatively, whether there exists a functional gradient of computational properties. Finally, the combination of a short processing time (< 1 min per 1000 neurons), a computational complexity that scales near-linearly with the size of the neuron population (Fig. 12), and ease of parallelization, makes SIMNETS an appealing tool for exploring very large-scale neuron populations.

### Algorithm Validation

Our analysis of simulated data with ground-truth subnetworks and known single neuron properties (i.e., encoded content and encoding timescale) demonstrates the current implementation of the SIMNETS algorithm can capture a neuron’s local and global computational relationships, even under conditions where computationally similar neurons relied on very different encoding schemes (e.g., rate coding vs. fine temporal coding schemes). This “agnosticism” can be beneficial when analyzing neurons that generate highly heterogeneous spike train outputs because of differences in their internal machinery (e.g., inhibitory vs. excitatory neurons) yet still reflect task-relevant information (Churchland and Shenoy, 2007; Diba et al., 2014; Martina et al., 1998). We demonstrated how the VP metric temporal accuracy parameter (q) could be used to interrogate the effect of temporal resolution on the computational configuration of the neurons.

The application of SIMNETS to three publicly available multi-neuron recording datasets validated the capabilities of the method in revealing known dominant feature representation in V1, M1, and CA1 (orientation, direction, place, respectively) without imposing stimulus or movement-driven parametric tuning models *a priori.* Of note, the results show that the computational properties of neurons are not uniformly distributed, but instead are organized into apparent subnetworks (i.e., CS clusters in the low-dimensional CS map) that emphasize certain collections of feature representations. Our results also suggest that SIMNETS may be able to detect distinct subnetwork structures hypothesized to support ensemble place-coding or complex feature conjunctions (Buzsáki, 2019; Diba et al., 2014). Although it was beyond the scope of this report to demonstrate the functional or computational significance of the detected putative subnetworks, our results suggest that the detected clusters are physiologically meaningful; our subsequent unsupervised cluster validation statistical procedure further provided support to these findings. Finally, our choice of datasets allowed us to demonstrate that this method generalizes well to neural recordings from a variety of brain regions, including sensory areas (Kohn and Smith, 2016), motor areas (Rao and Donoghue, 2014), and memory/cognitive areas (Pastalkova et al., 2008); different species, including rat and non-human primate; and across different recording technologies including laminar silicone probes (Buzsáki, 2004) and fixed-dept multi-electrode arrays (Maynard et al., 1997).

Collectively, our results demonstrate that single neuron SSIM matrices provide a simple, scalable, and powerful format for characterizing the computational properties of neurons, as well as identifying computational similarities and differences between them. In addition, examining the arrangement of neurons in the CS maps may suggest novel coding schemes that are yet unexplored by experimenters, generating new hypotheses for future experiments.

### Comparison to Existing Approaches

The SIMNETS analysis framework builds upon rich theoretical and mathematical literature spanning multiple domains and disciplines. Geometric and metric space representational models of similarity have a long history of application in the field of psychology where they have been used to model the perceptual relationships between sensory stimuli as latent or low-dimensional perceptual metric-space (Forstmann, 2011; Goldstone et al., 1991; Hyman, 1974; Shepard and Susan Chipman., 1970; Shepard, 1987, 1964; Thurstone, 2017; Tversky, 1988; Zenker and Gärdenfors, 2015). This approach has recently been adapted to study the intrinsic structure of neural representational spaces in visual, motor, and cognitive brain regions (for reviews, see; Edelman et al., 1998; Haxby, 2012; Kriegeskorte et al., 2008; Kriegeskorte and Kievit, 2013; Lehky et al., 2013). Several mathematical variations of this framework have been developed to study the information content of macroscale fMRI BOLD signals (Diedrichsen and Kriegeskorte, 2017; Haxby et al., 2001; Kriegeskorte and Bandettini, 2007), and both population-level and neuron-level spiking patterns (Houghton and Victor, 2010; Kiani et al., 2007; Vargas-Irwin et al., 2015a, 2015b; Victor and Purpura, 1996). To the best of our knowledge, this is the first work that uses a measure of 2nd-order point-process similarity analysis to generate latent space embedding of a collection of individual neurons.

The concept of a low-dimensional embedding that captures the functional relationship between spiking neurons was introduced in the seminal papers by (Baker and Gerstein, 2000; Gerstein and Michalski, 1981) describing the use of “Gravitational Clustering” (Wright, 1977): a neuron clustering and visualization tool for identifying groups of neurons with synchronous spiking patterns. This method is based on an analogy of the physics of the gravitational forces governing the dynamics and interactions of macroscopic particles. It treats the N neurons as N particles moving within an N-dimensional space, where charges that influence the attractive and repulsive interactions between particles are dictated by the temporal dynamics of pairwise synchronous spiking activity between neurons. The result is a visualization of particle clusters (and their trajectories) that represent dynamically evolving assemblies of synchronously active neurons. More recent work by Kiani et al. (2015) involves the application of the dimensionality reduction techniques (i.e., multidimensional scaling) to pairwise measures of between-neuron spike rate covariations to detect natural functional modules in a population of pre-arcuate and motor cortex neurons (Kiani et al., 2015). Although the goal of this method is like that of SIMNETS – to the extent that it makes use of single-trial information to group neurons in an unsupervised manner – this approach is applied to binned spike rates and appears to capture the covariation information carried in the absolute firing rates of the neurons’ responses across trials (Kiani et al., 2015), rather than the information carried in the neurons’ intrinsic spike train relational geometry. However, our analysis of the synthetic neurons using temporal codes, as well as rat CA1 neurons demonstrate that SIMNETS can detect a broader range of meaningful information processing motifs that reflect both condition-independent and condition-dependent information carried in the fine temporal structure of spike train outputs.

Yang et al. (2019) identified functional groupings of artificial neurons in a neural network by applying t-SNE to activation-based measures of single neuron selectivity across 20 different tasks (i.e., task variance) (Yang et al., 2019). The method is similar to the SIMNETS method in that it projects neurons onto a CS map according to the similarity of measures of output variance. This method requires knowledge of *trial labels* within tasks to group neurons according to the similarities of their changes in selectivity across tasks. By contrast, SIMNETS doesn’t require *a priori* knowledge about the similarities or differences of the information encoded on certain trials (i.e., trial labels), only that the trials are recorded simultaneously. This feature of SIMNETS is particularly valuable when trying to identify groups of functionally similar neurons in experiments with awake and freely moving animals, where the assumption of repeatable perceptual, cognitive, or behavioral states across trials is not always possible.

Several previous studies have used pairwise comparisons between the spike trains of different neurons to identify putative subnetworks or cell assemblies (Aertsen et al., 1989; De Blasi et al., 2019; Gerstein et al., 1985; Grün et al., 2002a; Humphries, 2011; Roudi et al., 2015; Singer and Gray, 1995). As with the gravitational clustering method, these studies have operated under the working hypothesis that the detection of time-series spike time or rate correlations is a signature of a potential functional link (Cole et al., 2016; Roudi et al., 2015; Singer and Gray, 1995; Ts’o et al., 1986). By contrast, SIMNETS identifies neurons with similar informational content even if they exhibit heterogeneous spike train outputs or utilize different encoding timescales (e.g., rate vs. precise spike timing). For the simulated neuron population, SIMNETS successfully identified clusters of the simulated neurons according to their ground-truth functional subnetworks and by increasing the sensitivity of the VP metric, the CS map was able to reveal further sub-groupings within the three identified functional CS clusters that highlighted subtle differences their output space geometries that corresponded to the different encoding timescale. This feature of SIMNETS could be particularly useful for determining if neurons encoded information across heterogenous timescales, e.g., fast vs. slow-timescale inhibitory neurons, or if they compute across a hierarchy of timescales (Cusinato et al., 2022; Fortenberry et al., 2012; Gorchetchnikov and Grossberg, 2007; Honey et al., 2012). Additionally, this feature of SIMNETS enabled us to cluster physiologically and functionally distinct groups of neurons in the CA1 dataset (i.e., place-cells vs non-place cells), which were not obvious when VP metric was insensitive to spike times (i.e., VP metric q = 0; data not shown).

Capturing the computational similarity between pairs of neurons in a more general way could be accomplished using Information theoretic approaches (Shannon, 1997), which could include estimating the shared or mutual information between the output spike trains of different neurons. Other approaches focus on the asymmetry of the predictive power between variables at different lags, resulting in “directed” estimates of functional connectivity such as Granger Causality, Transfer Entropy, or the Directed Transfer Function (Granger, 1983; Ito et al., 2011; Ursino et al., 2020). These strategies are based on estimating joint probability distributions across the activity patterns of pairs of neurons. A similar approach can be applied to relationships between multiple neurons using Generalized Linear Models (Chen et al., 2009; Truccolo et al., 2005). The number of possible activity patterns is very large, so this type of calculation can be challenging even when relatively large amounts of data are available. Using large amounts of data presents additional problems for this approach because it assumes that relationships between neurons remain relatively constant across the time span used to fit the models. Hence, every additional neuron requires exponentially more data and computing power and analyzing hundreds to thousands of simultaneously recorded neurons becomes intractable.

### Mitigating Experimenter Bias

A considerable amount of research in systems neuroscience has focused on identifying new classes of neurons based on their information-processing properties. The standard approach for many of these experiments involves recording single unit activity while a certain experimental variable of interest is manipulated (for example, providing systematic changes in stimulus features, or eliciting different behavioral responses, etc.)(Meyer et al., 2016). Standard statistical tests (ANOVA, etc.) are then used to determine if each neuron displays significant changes in firing rate across the experimental conditions. The percentage of significant neurons is usually reported as a functionally distinct “class” of neurons sensitive to the variable of interest. It is common to exclude neurons that do not reach statistical significance or cannot be fit using a predetermined model from further analysis. This approach is prone to both selection and confirmation bias, and ultimately produces “classes” of neurons identified based on arbitrary statistical thresholds imposed on what are likely continuous distributions of properties. The SIMNETS analysis framework is an unsupervised approach to determine if neurons are organized across a functional continuum or are organized into statistically separable functional classes, thereby mitigating the experimenter’s bias inherent in parametric neural discovery methods. In addition to providing a principled way to determine if a consistent organization of information processing modules can be found across sessions and subjects, we believe that the ability to intuitively visualize relationships within networks of neurons will provide a unique perspective leading to new data-driven hypotheses and experimental refinement.

### Limitations & Considerations

Several important limitations of SIMNETS are worth noting. First, estimates of similarity using spike train metrics require that the time windows of interest be of equal length, making it difficult to compare neural responses with different time courses. This weakness is common to all trial-averaging models commonly used in the literature that we are aware of.

Second, although the SIMNETS framework does not require a priori assumptions about the variables potentially encoded by neural activity, experimental design and data selection will still have a direct effect on the results obtained. For example, a set of neurons identified as a functional subnetwork could separate into smaller groups with different computational properties when additional task conditions are added to the analysis. Thus, the functional properties identified using SIMNETS are only valid within the context of the data examined and may not necessarily extrapolate to different experimental conditions. Ultimately, the computational space of the population is only as rich as the experimental design allows (Gao and Ganguli, 2015).

Third, SIMNETS does not provide a means to assess sources of variance. For example, subsets of neurons may exhibit systematic changes in spike train outputs through an experimental session, reward experience, timing, attention, or many other variables which can be revealed by the neural data; however, it will be up to the experimenter to identify their source. Nevertheless, SIMNETS can guide hypothesis generation for experiments that identify these influences.

Fourth, it is possible that neuronal subnetworks are constantly re-arranged depending on changing external or internal signals, e.g., ethological demands, task demands, or task stages, or attentional levels. For example, a neuron could potentially exhibit rapid changes in its computational/functional interrelationships with other neurons during processing epochs that are shorter than the selected analysis time window. The current version of the SIMNETS algorithm was not designed to distinguish between rapidly changing sub-network memberships. However, applying the SIMNETS algorithm multiple times over different epochs can help with determining if different network configurations are engaged at different times.

#### Contributions

Using ordering of author list. Conceptualization: all authors; Methodology: JH, DB, JZ, CVI. Software: JH, DB, JZ, CVI. Formal Analysis: JH. Dataset Curation: JH, CVI. Figures: JH. Writing, Review & Editing: all authors. Writing, Manuscript Preparation: JH. Supervision: CVI, JPD. Funding Acquisition: CVI, JPD.

## Acknowledgments

This work was supported by NIH Director’s New Innovator award (1DP2NS111817-01), NINDS-Javits (NS25074), The Israeli Brain Prize, and the Killam Trust Award Foundation. We thank Stuart Geman and Adam Kohn for their feedback on the general method, as well as M. Nevor and J. Murphy for their assistance with animal care and instrumentation design, and N. Tolley developing a python-based code repository. We also thank the Buzsáki and Kohn Lab for allowing us (and the field) to work with their data via the CRCNS data repository. The views expressed in this article are those of the authors and do not necessarily reflect the position or policy of the NIH, the Department of Veterans Affairs, or the United States government.

## Code and Data Availability

The SIMNETS software is a C++ optimized MATLAB Package available on GitHub at: https://donoghuelab.github.io/SIMNETS-Analysis-Toolbox/. This includes an interactive MATLAB Live tutorial (.mlx).

## Supplementary Information List

Supplementary Fig. S1. Choice of dimensionality reduction method impacts the structure of the low-dimensional CS neuron map.

Supplementary Fig. S2. Stability of cluster evaluation as a function of different SIMNETS parameters and CS map extrinsic dimensionality number.

Supplementary Fig. S3. Supplementary CA1 Hippocampal analysis provides extended view SIMNETS and SSIMS maps (cont. from Fig. 11)

## Supplementary Information

**Fig. S1.**
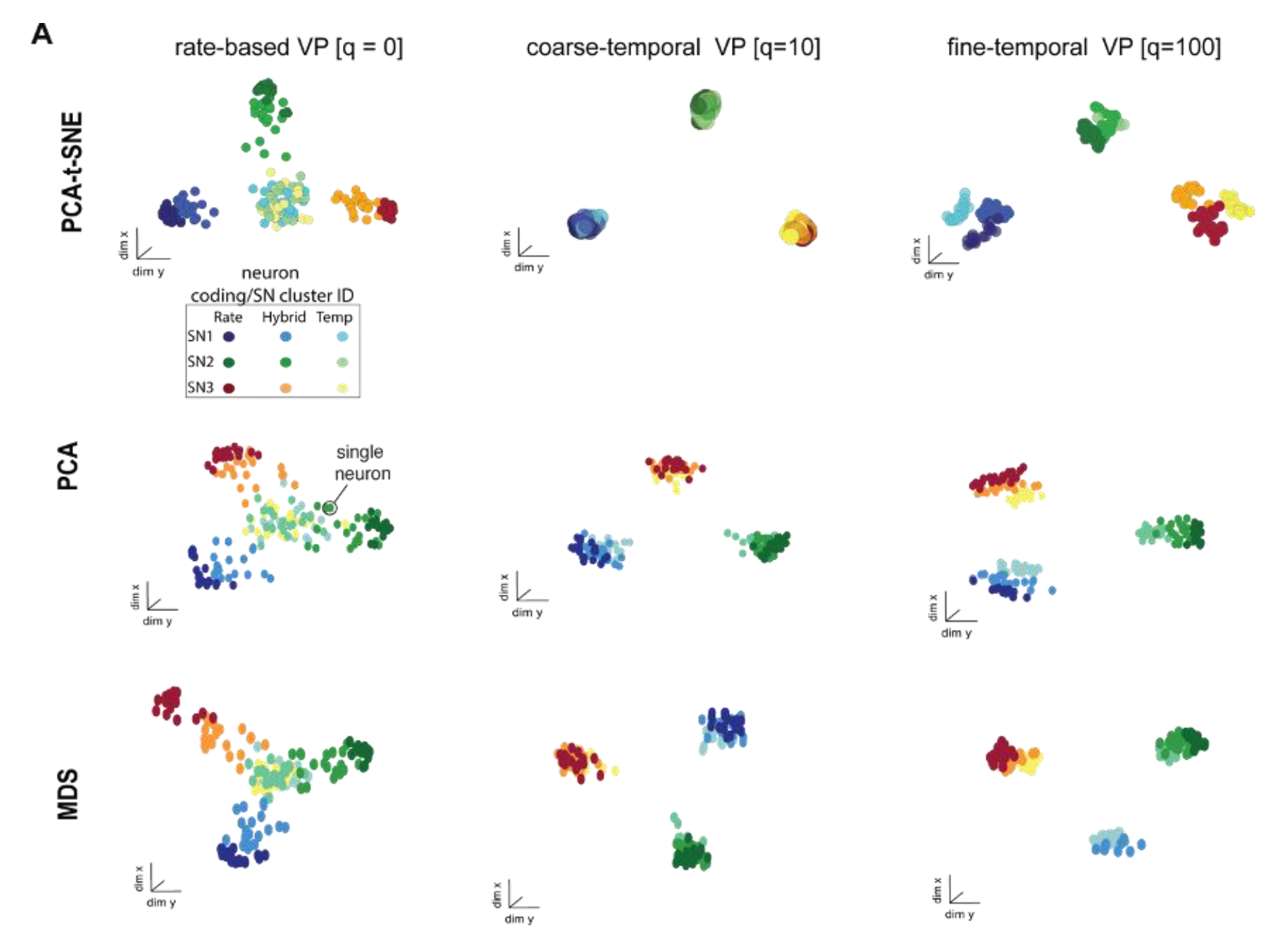
Choice of dimensionality reduction method impacts the structure of the low-dimensional CS neuron map. **A)** Columns: low-dimensional CS neuron mapping of simulated neuron population dataset for three different VP temporal accuracy values (same as Fig. 5). Rows: the same CS neuron map using PCA-initialized t-SNE (PCA-tSNE) (top row), PCA (middle row), and multidimensional Scaling (MDS, bottom row) (Kruskal and Wish, 1978). Individual points represent individual simulated neurons, and the color labels indicate their simulated ground-truth properties, including coding type and ground-truth CS neuron cluster. PCA is a linear global method; MDS is a local linear method, while PCA-initialized t-SNE combines both linear/global and non-linear/local techniques.

**Fig. S2.**
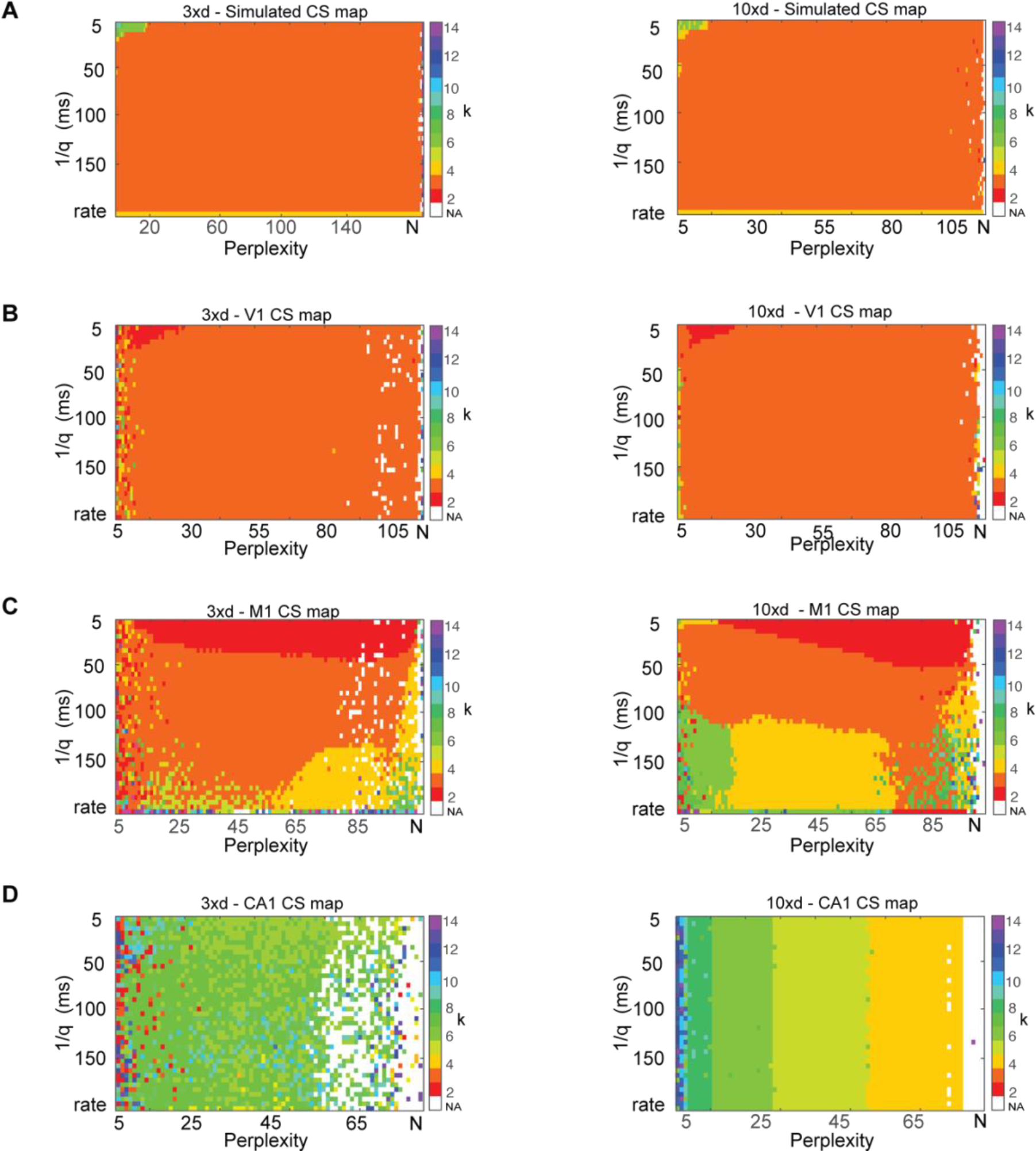
Stability of cluster evaluation as a function of different SIMNETS parameters and CS map extrinsic dimensionality number. SIMNETS clusters (*k*) as function of *perplexity* and temporal accuracy (1/*q*) in a 3xN PCA-tSNE projection space (*left*) and a 10-*d* PCA-tSNE projection space (*right*) for the simulated neuron population (N =180, neuron’s) (**A**), the V1 neuron population (**B**), the M1 neuron population (**C**), and the Hippocampal CA1 neuron population (**D**). Color bars indicate the optimal number of natural clusters (optimal k-means partitions) within the CS neuron maps, which was determined using a silhouette analysis.

**Fig. S3.**
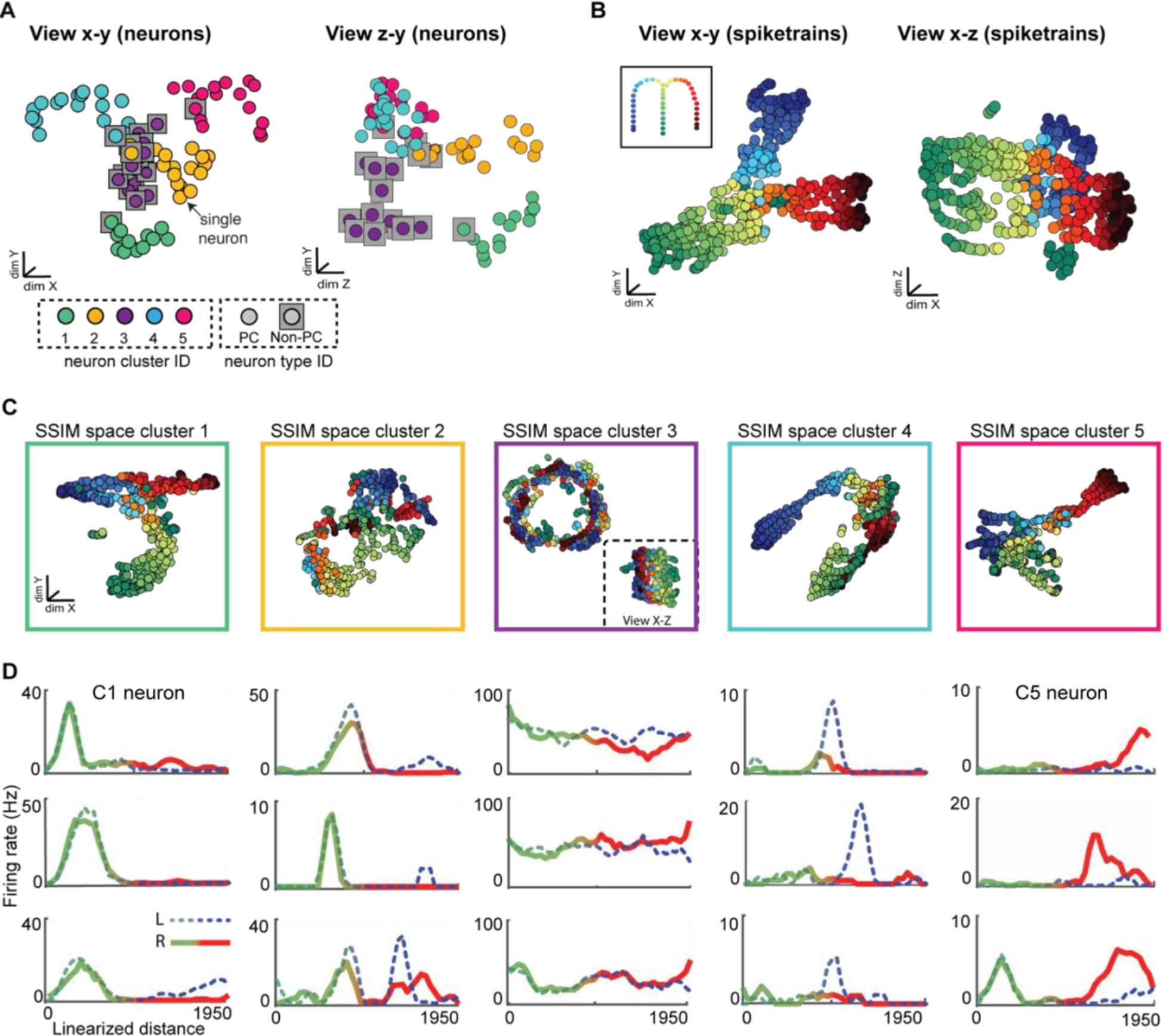
Supplementary CA1 Hippocampal analysis provides extended view SIMNETS and SSIMS maps (cont. from Fig. 11). **A)** CA1 population SIMNETS CS map from different viewing angles. *Left*: x-y viewing angle of CS map (same as **Fig. 11A**); *Right*: z-y viewing angle. Points are neurons; gray squares circumscribe those neurons categorized as non-PC neurons. Color corresponds to the k-means CS neuron clusters. **B)** *Left*: X-Y view of population-level SSIM map. Points are individual spike trains and colors indicate the position in the maze. *Left*: (same as Fig. 11D). *Right*: x-z viewing angle. **C)** Cluster-level SSIMS maps for all five CS neuron clusters (cluster c3-c5 same as **Fig. 11B**). Outer colored squares correspond to the neuron CS cluster labels in (**A**). The inset cluster c3 neuron (*purple*) provides an additional x-z viewing for the cluster c3 neuron (*purple*) to better highlight its complex shape. **D)** Each column shows three representative single neuron trial-averaged firing-rate functions for the CS neuron clusters c1-c5, respectively. The x-axis corresponds to the linearized distance traveled by the rat along a given arm of the track. The broken green-blue trace corresponds to a left (L) maze run and the solid green-red trace corresponds to right (R) maze run. Rate functions resulted from binning and averaging firing rates across left or right trials and smoothing over with a sliding window.

## Supplementary Methods

### Cluster Validation: Shuffle-based Significance Test Silhouette Analysis

Our CS neuron clusters validation process consists of using a silhouette graphical analysis to determine the natural number of clusters within the empirical CS neuron map (Rousseeuw, 1987) and our novel shuffle-based statistical procedure to verify the statistical validity of these results. Silhouette analysis is used to assess the quality of the clustering obtained from different partitions of a given dataset (in this case, obtained using different values of k for the k-means algorithm). An optimal number of clusters is selected as the partition number, k, that maximizes the average silhouette value, referred to as the Silhouette Coefficient (SC). A silhouette value, h_i_, is a measure of how close a point is to other data points in its assigned cluster as compared to points in other clusters:

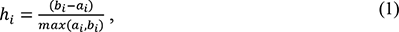

where a_i_ is the average distance between each point within a given cluster and all other points within that same cluster, and b_i_ is the average distance to each point and all points in other clusters, minimized over all possible cluster configurations. Silhouette values are typically normalized to ([0 to 1]), where a high value indicates that a given point is well matched to its own cluster and far from neighboring clusters. In general, a maximized average silhouette below 0.25 indicates data that are not structured while a value below 0.5 would indicate poor or potentially spurious clusters. In the next section, we outline a procedure for testing the statistical significance of the optimal cluster number identified using the Silhouette analysis.

We developed a significance test for the purpose of determining the likelihood of detecting a given number of clusters by chance using silhouette analysis under the null hypothesis that there is no genuine covariation relationship between the inherent structures of the SSIM matrices. The significance test involves generating a null distribution of silhouette values based on shuffled data across a range of partition (k) values. In SIMNETS, computational similarities are captured by the pairwise measures of correlation between each neuron’s respective spike train distances values in the single neuron SSIM matrices. Our test relies on a shuffling procedure that destroys the pairwise dependencies between the SSIM matrices, and subsequently, any significant measures of correlation in the CS neuron matrix. This approach is inspired by the Mantel test, a permutation-based procedure that tests the significance of the observed correlation between two symmetric matrices (Legendre, 2000; Mantel, 1968). The intuition of a Mantel test is that if a significant relationship exists between the values of matrix A and matrix B, then randomizing the rows and columns of one matrix will destroy any existing dependencies. As a result, the correlation between the shuffled matrix pair will tend to be lower than the original correlation value observed between the un-shuffled matrix pair. The probability of observing r(A, B) is then calculated as the proportion of permutations for which the shuffled correlation measures are smaller than or equal to r(A, B). Here, we carry out a similar permutation operation on the SSIM matrices, in that we destroy any dependencies that exist between the matrices; however, we use the average silhouette value as the test statistic, rather than the correlation values, as is the case with the Mantel test.

The procedure involves symmetrically shuffling the rows/columns of each N SSIM matrix separately and re-calculating the pairwise correlations between the SSIM matrices to generate a new NxN CS matrix. This NxN CS matrix is then transformed into a new CS map using t-SNE, and a new set of silhouette values is calculated for the range of tested partition values. This procedure is repeated to generate a null distribution of average silhouette values that are approximately normally distributed (e.g., 1000+ iterations). If the observed maximized average silhouette value, SC, falls above the empirically calculated (1-α)100% confidence interval, then the detected number of clusters is considered statistically meaningful.

**Table 1:**
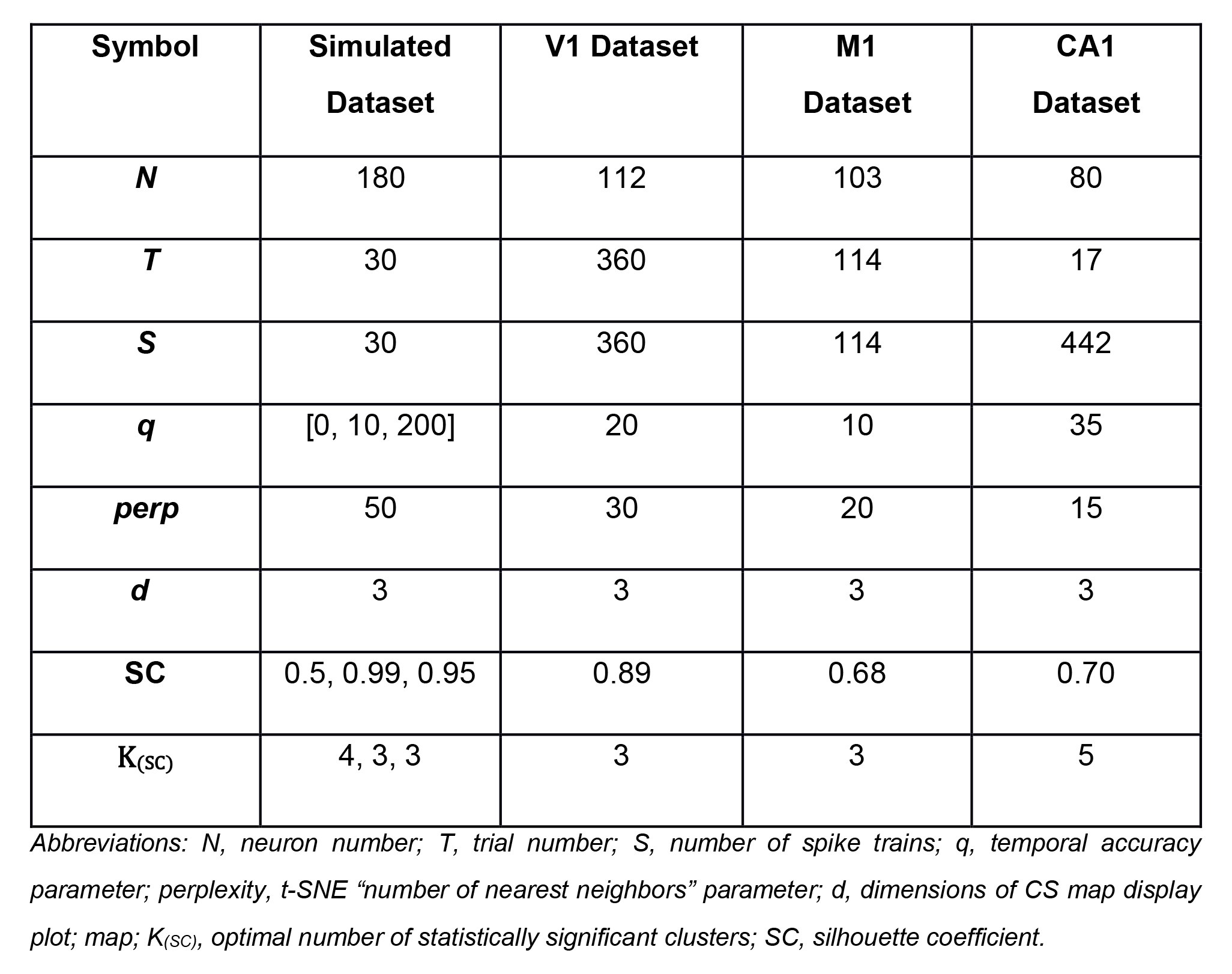
Summary of SIMNETS algorithm Inputs/Outputs across datasets.

| Symbol | Simulated Dataset | V1 Dataset | M1 Dataset | CA1 Dataset |
| --- | --- | --- | --- | --- |
| $N$ | 180 | 112 | 103 | 80 |
| $T$ | 30 | 360 | 114 | 17 |
| $S$ | 30 | 360 | 114 | 442 |
| $q$ | [0, 10, 200] | 20 | 10 | 35 |
| $perp$ | 50 | 30 | 20 | 15 |
| $d$ | 3 | 3 | 3 | 3 |
| $SC$ | 0.5, 0.99, 0.95 | 0.89 | 0.68 | 0.70 |
| $K_{(SC)}$ | 4, 3, 3 | 3 | 3 | 5 |

Abbreviations: $N$ , neuron number; $T$ , trial number; $S$ , number of spike trains; $q$ , temporal accuracy parameter; $perp$ , perplexity, $t$ -SNE “number of nearest neighbors” parameter; $d$ , dimensions of CS map display plot; $K_{(SC)}$ , optimal number of statistically significant clusters; $SC$ , silhouette coefficient.

#### Code and Data Availability

C++ optimized MATLAB code and tutorial are available on GitHub at: https://donoghuelab.github.io/SIMNETS-Analysis-Toolbox/

### Simulated Dataset – Data Simulation and Analysis

Spike train Simulation We simulated the spiking activity of a population of N = 180 synthetic neurons that consisted of 3 computationally distinct subnetworks (SN1, SN2, SN3) of 60 neurons. Each subnetwork was designed to produce similar spike trains for two non-modulating conditions, referred to as the “baseline” conditions, and a different pattern for a third condition, referred to as the “modulating” condition. For example, subnetwork SN1 was modulated during condition A and exhibited the same baseline activity spike pattern during both conditions B and C, whereas subnetwork SN1 was modulated during condition B and exhibited the same baseline spike pattern during conditions A and C, etc. The neurons within each subnetwork could be further divided into three sub-groups of n = 20 neurons, where each sub-group altered their spike-train patterns between the active and baseline states according to one of three different encoding strategies:

i. Rate coding: firing rate increased by 50% for the modulating condition (all spike times were randomly chosen)
ii. Temporal coding: the two baseline conditions and the modulating condition were associated with specific (randomly generated) temporal sequences of spikes. The number of spikes was kept constant across baseline and modulating conditions. Spike times were jittered by +/- 5 ms for each trial.
iii. Mixed temporal/rate coding: similar to the temporal coding, but additionally, the modulating condition included 25% more spikes.

To simulate stochastic variation in spiking patterns, 50% of the spikes were randomly removed for each condition. A total of 30 seconds of simulated recording time was generated, with the trial condition changing every second between A, B, and C patterns. This example dataset is included in the SIMNETS software package included with this submission. For MATLAB Tutorial, see:

https://donoghuelab.github.io/SIMNETS-Analysis-Toolbox/SIMNETS.mlx

### Neural Datasets – Task Description and Data Analysis

#### Primate Primary Visual Cortex Dataset

##### Task Description

We analyzed a previously described dataset of 112 primary visual (V1) single-units (which we refer to as neurons) recorded in an anesthetized Macaca Fascicularis using a 96-channel microelectrode array (Kohn and Smith, 2016; Smith and Kohn, 2008). We analyzed the data from a single subject (monkey 3, single session). Briefly, sinusoidal gratings were presented at 6 different orientations *θ* = {0°, 30°, 60°, 90°, 120°, 150°} and 2 drift directions (rightward and leftward drift, orthogonal to orientation). Each stimulus was presented 112 times for 1.28 seconds. The position and size of the stimuli was sufficient to cover the receptive fields of all recorded neurons. For more details on the task design and data pre-processing, *see* Kohn and Smith, 2016 and Smith and Kohn, 2008, or go to the CRCNS.org data repository: https://crcns.org/data-sets/vc/pvc-11/about).

##### Single Neuron Tuning Analysis

The single neuron analyses described were applied to the same set of spike trains used in the SIMNETS analysis. This involved extracting 1 second of spiking data from the first 30 repetitions of each stimulus (S = 360, spike trains), starting 0.28 seconds after stimulus onset. Only a small fraction of the total number of recorded trials was used in the analysis (25%) as we wanted to demonstrate SIMNETS ability to cluster neurons in datasets where only a small number of trials are available. These analyses were carried out independently from the SIMNETS analysis and were not used to guide the SIMNETS algorithm in any way.

We characterized the preferred orientation of each V1 neuron by fitting a Gaussian distribution to the firing rate function R:

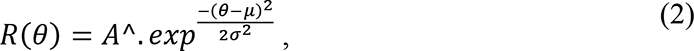

where is the stimulus orientation, Â is the peak response, μ the mean, and σ^2^ is the variance of the Gaussian. The function takes on a maximum value at 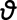 = μ, for θ = [0, 180), which corresponds to the neuron’s preferred orientation. Drift-direction was not explicitly repotted in this work, but it was taken into consideration by calculating preferred orientation separately for the different drift directions (i.e., 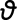 = [0, 180) and 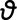 = [180, 360)). Summary statistics of drift direction were not reported but an example neuron with direction-of-motion dependent orientation selectivity is presented in the main text (Fig. 7C, see example n36 in CS cluster 3). We calculated the normalized “peak-to-trough” of the orientation response function, the orientation index (OI), as an indicator of orientation tuning strength:

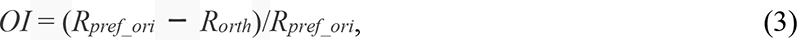

where *R_pref_ori_* is the “peak response” of the orientation tuning function (See: eqt. 2, equivalent to R(θ), and *R_orth_* is the “trough response” at the orthogonal angle (or function minimum) of the tuning function (Mazurek et al., 2014).

##### V1 SIMNETS CS Map Characterization

We used a circular-linear correlation (r_cl_) analysis to assess SIMNETS’ ability to organize neurons according to their computational properties (Berens, 2009). The correlation between each neuron’s preferred orientation and its location along each dimension in the low dimensional map y_i_ was calculated using:

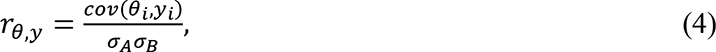

, where *σA* and *σB* are the standard deviations of the neurons’ preferred orientations and y represents the neurons’ locations in the map. A high correlation value indicates a strong relationship between a neuron’s preferred orientation/direction and map location and demonstrates that functionally similar neurons were mapped to nearby regions of the map. The r_cl_ value for the dimension with the highest value was reported.

#### Primate Primary Motor Cortex Dataset

**Task Description.** SIMNETS was applied to the previously described dataset of *Macaca mulatta* primary motor (M1) cortex neurons (i.e., single-units) recorded during a planar 8- direction reaching task (Rao and Donoghue, 2014). The single-unit activity was simultaneously recorded from the upper limb area of the primary motor cortex using a chronically implanted microelectrode array. The monkey was operantly trained to move a cursor that matched its hand location to targets projected onto a horizontal reflective surface. A visual cue was used to signal movement direction during a variable duration instructed delay period (1 – 1.6 s) to one of eight radially distributed targets on the screen with the associated reach angles of *φ* = {0°, 45°, 90°, 135°, 180°, 225°, 270°, 315°}. At the end of the instructed delay period, the central target was extinguished, instructing the monkey to reach towards the previously cued target.

##### Single Neuron Tuning Analysis

We analyzed 1 second of neural data from correct trials (S = 114), starting 0.1 second before movement onset. Characterization of the detected SIMNETS clusters is like that described in the previous section. We characterized the preferred movement direction of each M1 neuron by fitting a von Mises distribution to the firing rate function R:

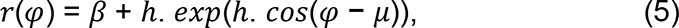

where *β* is the offset of the function, h is the depth of the tuning, is the reach angle and μ is preferred reach direction of the cell (Mardia and Zemroch, 1975). The function takes on a maximum value at μ, which corresponds to the neuron’s preferred direction. The tuning functions were then used to calculate each neuron’s tuning depth, or the reach direction index (RDI). This was calculated as the normalized peak-to-trough responses using an equation of similar form to the V1 neuron OI (See Equation (3)).

#### Rat Hippocampal CA1 Dataset

##### Task Description

We applied SIMNETS to a previously described dataset of rat hippocampal neurons made publicly available by the Collaborative Research in Computational Neuroscience (CRCNS.ORG) data-sharing repository (Pastalkova et al., 2015, 2008). We analyzed the data from a single subject rat during one session (dataset: hc-5 01_maze06_MS.002). The neurons were simultaneously recorded from the CA1 hippocampal region using multi-site silicon probes while the rat performed a spatial navigation task in a maze. Briefly, the rat was trained to run through the arms of a “figure-8” maze in a left/right alternating manner to receive a reward. The left/right track runs were interleaved with a wheel-run period that functionally served as a memory delay-period. The rat performed T = 17 correct trials (Right= 8, left trials; T*_left_*= 9, right trials), taking on average 4.3 seconds to reach the rewards located at either end of the arms. The rat’s path along each arm of the track was linearized and divided into small spatial bins (80cm) for the SIMNETS analysis, respectively. For additional details go to CRCNS.org repository: https://crcns.org/data-sets/hc/hc-5/about-hc-5).

##### Single Neuron Place Field Analysis

The rat’s path along each arm of the track was linearized and divided into 80 mm spatial bins when generating the spatial firing field maps. Bins corresponding to reward locations and the inter-trial activity were excluded from the analysis, leaving a total of 390 bins for each of the left and right trajectories, where the first 19 spatial bins were common to both trajectories. We generated a separate spatial firing map for the left and right trajectories of each neuron by dividing the number of spikes in the i-th bin by the rat’s occupancy time t_i_ and used a Gaussian kernel (width = 3 bins/150 mm) to smooth across the firing rates in each bin. Neurons that did not exhibit a 5 Hz firing rate in at least 1 spatial bin were not included in the analysis, leaving a total of N = 80 neurons. We characterized the neurons as non-place cells (n = 20, non-PC) or place cells (n = 60, PC) based on their spatial firing properties and an information-theoretic measure of the spatial information in their spikes (Eqt. 6) (Skaggs et al., 1992a). Neurons were classified as having place cell-like activity if the firing rate in three contiguous bins exceeded the mean of all other firing fields by 20% (Jeffery et al., 1997; Skaggs et al., 1992)(using 2.5 STD of the out-of-field firing rate produced similar results) and if their information content exceeded 0.5 bits/spike on either the left or right trajectories (Skaggs et al., 1992). The spatial information metric, I_spike_, is a measure of the extent to which a neuron’s spiking activity can be used to predict the rat’s position along the track. The spatial information content of the neuron (measured in bits/spike) is defined as:

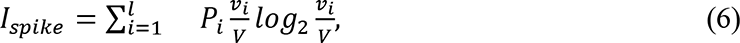

where Pi is the occupancy probability, *νi* is the firing rate in the i-th bin, and V is the overall mean firing rate of the cell across all bins in trajectory.

